# Growth inhibitory factor/metallothionein-3 is a sulfane sulfur-binding protein

**DOI:** 10.1101/2023.10.19.563042

**Authors:** Yasuhiro Shinkai, Yunjie Ding, Toru Matsui, George Devitt, Masahiro Akiyama, Tang-Long Shen, Motohiro Nishida, Tomoaki Ida, Takaaki Akaike, Sumeet Mahajan, Jon M. Fukuto, Yasuteru Shigeta, Yoshito Kumagai

## Abstract

Cysteine-bound sulfane sulfur atoms in proteins have received much attention as key factors in cellular redox homeostasis. However, the role of sulfane sulfur in zinc regulation has been underinvestigated. We report here that cysteine-bound sulfane sulfur atoms serve as ligands to hold and release zinc ions in growth inhibitory factor (GIF)/metallothionein-3 (MT-3) with an unexpected C–S–S–Zn structure. Oxidation of such a zinc/persulfide cluster in Zn_7_GIF/MT-3 results in the release of zinc ions, and intramolecular tetrasulfide bridges in apo-GIF/MT-3 efficiently undergo S–S bond cleavage by thioredoxin to regenerate Zn_7_GIF/MT-3. Three-dimensional molecular modeling confirmed the critical role of the persulfide group in the thermostability and Zn-binding affinity of GIF/MT-3. The present discovery raises the fascinating possibility that the function of other Zn-binding proteins is controlled by sulfane sulfur.

## Introduction

Sulfane sulfur is a chemical state of the sulfur atom with six valence electrons that are covalently bound to sulfur atoms (*1, 2*). Growing evidence supports the widespread existence of hydropersulfidated and polysulfidated proteins in all cell types, referred to as sulfane sulfur-binding proteins (SSBPs) (*3*). Protein sulfuration occurs via post-translational and co-translational pathways. Rhodanese is known to catalyze the production of sulfane sulfur and attach it to the thiol group of the protein itself using thiosulfate as a substrate (*4*). 3-Mercaptopyruvate sulfurtransferase is a rhodanese-like enzyme that uses 3-mercaptopyruvate as the preferred sulfur donor (*5*). Cystathionine β-synthase and cystathionine γ-lyase use cystine as a substrate and catalyze the production of sulfane sulfur-containing cysteine hydropersulfide (CysSSH)(*6*), whose terminal sulfane sulfur can be reversibly transferred to other thiols such as glutathione (GSH) or protein-SH to form GSH hydropersulfide (GSSH) or protein hydropersulfides (protein-SSH), respectively (*7*). Cysteinyl-tRNA synthetase 2 catalyzes the production of CysSSH from CysSH (*8*), thereby producing CysSSH-integrated nascent proteins. The biological function of sulfane sulfur has received considerable attention in redox biology because of its antioxidant/anti-electrophilic capacity. However, the role of sulfane sulfur in proteins is not fully understood; therefore, further mechanistic investigation is required. Although several SSBPs have been identified using various methods (*6, 9*), in the present study we used β-(4-hydroxyphenyl)ethyl iodoacetamide (HPE-IAM) (*8*) to derivatize the sulfane sulfur because we herein found that HPE-IAM has ability to extract sulfane sulfur atoms from SSBPs to form bis-S-HPE-AM adduct at a certain condition, thereby enabling quantitative analysis using LC-MS/MS with a stable isotope-labeled standard, bis-S^34^-HPE-AM.

In biological systems, cysteine-rich proteins can act as “redox switches” which sense accumulated oxidative stressors and free zinc ions, store excess metals, control the activity of metalloproteins, and serve as triggers for the activation of cellular redox signaling cascades (*10*). Metallothionein (MT), discovered in 1957 (*11*), is an important cysteine-rich metal-binding protein involved in three major biological processes: homeostasis of essential metals, detoxification of toxic metals, and protection from oxidative stress (*12, 13*). It is recognized that metal binding to MT is thermodynamically stable, but oxidation of the thiolate cluster readily leads to metal release and formation of intramolecular MT-disulfide linkages. Zinc ions released from zinc/thiolate clusters of MT are suggested to function as signaling molecules for cellular redox homeostasis (*14*). Simultaneously, reduction of MT-disulfide by cellular reducing agents can occur in a process called the “MT redox cycle” (*12*). However, the biochemical features of MT related to these functions have not been fully characterized. In addition, although the gas chromatography–flame photometric detector technique showed that MT isoforms contain sulfide ions (*15, 16*), it remains unclear if these sulfides are indeed sulfane sulfur atoms that act as essential factors in controlling protein redox states, thereby regulating cellular zinc homeostasis. Because of its constitutive expression, this study focused on MT-3, which was originally identified as a growth inhibitory factor (GIF) in the human brain (*17*). This study aimed to clarify the existence and content of sulfane sulfur in GIF/MT-3, the redox regulation of sulfane sulfur in holo- and apo-GIF/MT-3 in association with zinc release, and the effect of sulfane sulfur on the thermostability and metal binding affinity of GIF/MT-3. We found that sulfane sulfur atoms provide a redox-dependent switching mechanism for zinc/persulfide cluster formation in GIF/MT-3.

## Results

### Existence of persulfide and polysulfide groups in apo-GIF/MT-3

It is well recognized that GIF/MT-3 is able to bind seven zinc ions. To determine if GIF/MT-3 is an SSBP, we used an *Escherichia coli* expression system to prepare recombinant human Zn_7_GIF/MT-3 protein, from which apo-GIF/MT-3 was subsequently prepared (Fig. 1A). To detect sulfur modification in GIF/MT-3, we attempted to first measure the molecular weight of the whole protein with and without bound zinc. Using Fourier transform ion cyclotron resonance (FT-ICR)-MALDI-TOF/MS, Zn_7_GIF/MT-3 was mainly detected at *m*/*z* = 7071 (Fig. 1B), which corresponded to the mass of zinc-free apo-GIF/MT-3 and indicated that zinc dissociates from protein in the acidic conditions used for MALDI sample preparation. However, apo-GIF/MT-3 showed several peaks at *m*/*z* = 7071, 7085, 7103, and 7117, with a main peak at *m*/*z* = 7053 (Fig. 1C) that corresponded to oxidized GIF/MT-3 with nine intramolecular cysteine disulfide bonds, which presumably release the molecular weight equivalent of nine molecules of H_2_ (Fig. 1D). Thus, the peaks at *m*/*z* = 7085 (oxidized GIF/MT-3 plus one sulfur atom), 7103 (apo-GIF/MT-3 plus one sulfur atom), and 7117 (oxidized GIF/MT-3 plus two sulfur atoms) suggested that oxidized apo-GIF/MT-3 contains sulfane sulfur-like species that presumably exist as intramolecular cysteine trisulfide or tetrasulfide bridges (Fig. 1D). Note that an increase in mass of 32 Da can also result from addition of two oxygen atoms.

**Fig. 1.**
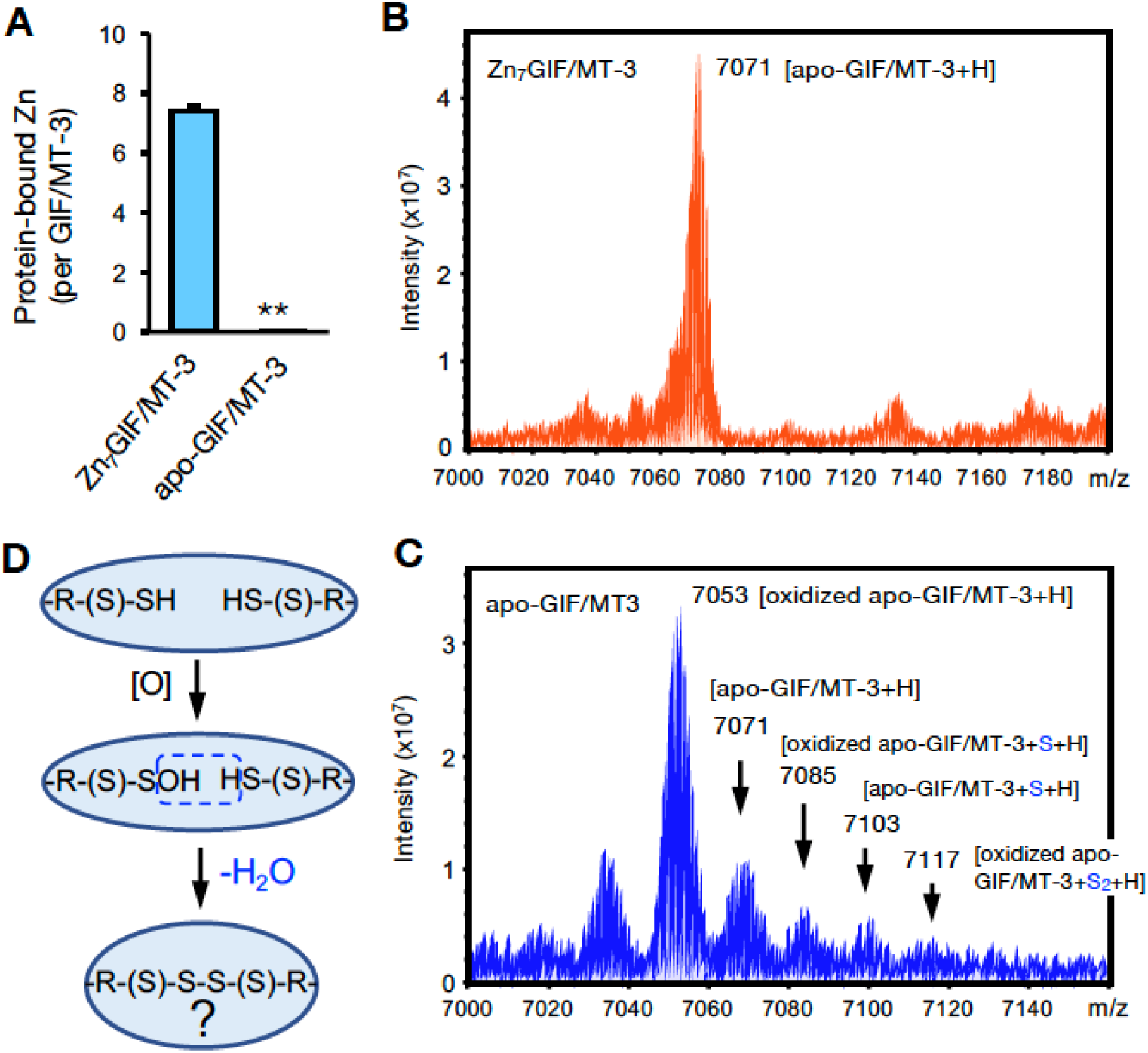
Detection of sulfane sulfur in GIF/MT-3 by MALDI-TOF/MS. (A) Preparation of recombinant Zn_7_GIF/MT-3 and oxidized apo-GIF/MT-3 proteins. Recombinant human Zn_7_GIF/MT-3 (10 μM) was incubated in HCl (0.1 N) at 37°C for 30 min and then replaced with 20 mM Tris-HCl (pH 7.5) buffer and incubated for 36 h at 37°C. After removal of low-molecular-weight molecules using 3 kDa centrifugal filtration, GIF/MT-3-bound zinc content was measured using ICP-MS. (B) FT-ICR-MALDI-TOF/MS spectrum (positive-ion mode) of Zn_7_GIF/MT-3. (C) FT-ICR-MALDI-TOF/MS spectrum (positive-ion mode) of oxidized apo-GIF/MT-3. (D) Putative oxidation reaction scheme in apo-GIF/MT-3 protein.

Raman spectroscopy is used to detect bonding changes in proteins, including MTs (*18, 19*). The Raman shift of Zn_7_GIF/MT-3 (Fig. 2A) contained a peak at 307 cm^−1^, which is attributable to both S-terminal and S-bridging ligands (e.g., S–Zn, S–Zn–S) (*18*). Also, the peaks at 761 cm^−1^ and 778 cm^−1^ presumably corresponded to cysteine–metal bonds (e.g., C–S–Zn) and backbone vibrations, respectively (*18*). Overall loss of zinc was indicated by the decrease in the intensity of these peaks in the Raman spectra of apo-GIF/MT-3 and GIF/MT-3 treated with HPE-IAM to consume sulfane sulfur atoms (Fig. 2A) and also confirmed using inductively-coupled plasma (ICP)-MS (Fig. 1A). The peak around 400 cm^−1^ is reported to correspond to vibrations of metal–S bridges (e.g., Zn–S–Zn) (*18*); however, loss of such a peak was not observed for apo-GIF/MT-3 and HPE-IAM-treated GIF/MT-3. The Raman peaks within the 510–520 cm^−1^ range reportedly indicate disulfide bonds in MT (*18*); in our spectra, an intense peak at 511 cm^−1^ was observed for apo- GIF/MT-3 but not Zn_7_GIF/MT-3 and HPE-IAM-treated GIF/MT-3 (Fig. 2A).

**Fig. 2.**
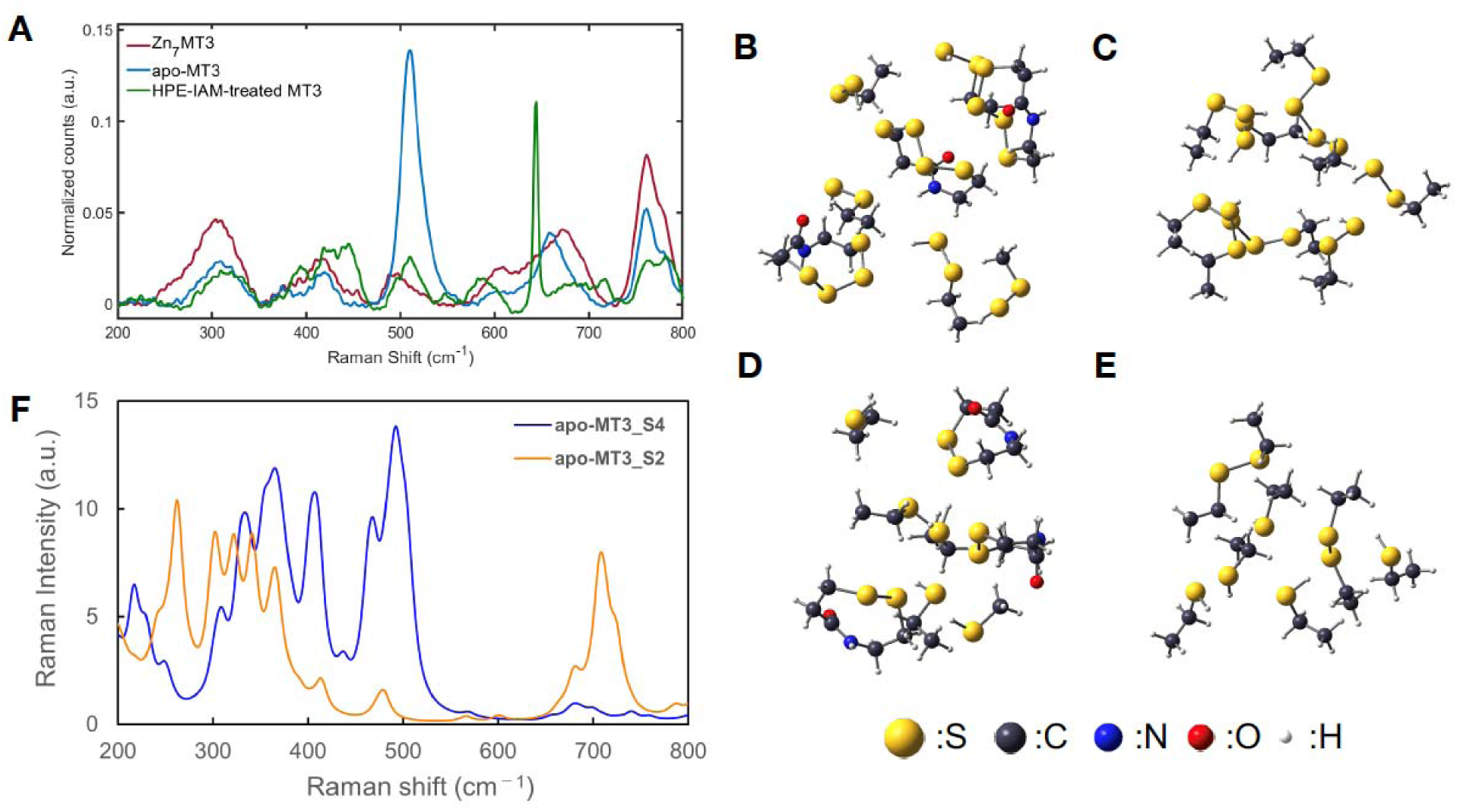
Detection of sulfane sulfur in GIF/MT-3 by Raman spectroscopy. (A) Raman spectra of Zn_7_GIF/MT-3, oxidized apo-GIF/MT-3, and HPE-IAM-treated GIF/MT-3 in the 250–800 cm^−1^ region. FT-ICR, Fourier transform ion cyclotron resonance. Optimized geometries for (B) α-domain and (C) β-domain models of apo-GIF/MT-3 (assuming some cysteines with persulfide and tetrasulfide bonds as shown). Optimized geometries for (D) α-domain and (E) β-domain models of apo-GIF/MT-3 (assuming some cysteines with disulfide bonds). (F) Calculated Raman spectra of apo-GIF/MT-3 models with/without sulfane sulfurs.

In general, the S–S bonds of polysulfides are strong Raman scatterers due to the high polarizability of the bonding and lone pair electrons at the two-coordinate sulfur atoms (*20*). It was also shown that S–S, S–S–S, and S–S–S–S structures exhibit different Raman shifts (*21, 22*) and the different Raman bands reported for polysulfide may correspond to different geometrical forms of the molecule. To confirm the origin of the Raman peak at 511 cm^−1^, quantum chemical calculations were made based on three-dimensional (3D) homology modeling, as described later, of apo-GIF/MT-3 structures with and without persulfides; the free cysteines and their persulfides were assumed to be protonated and some of the neighboring cysteines formed disulfides or tetrasulfides, depending on the model. The details of the model structures (Fig. 2B–E) and the calculation schemes are described in the Experimental procedures section. Figure 1I shows the calculated Raman spectra of apo-GIF/MT-3 with thiol (–SH) groups and disulfide (S–S) bonds (apo-GIF/MT-3_S2 model), and of apo-GIF/MT-3 persulfide (–SSH) groups and tetrasulfide (S–S–S–S) bonds (apo-GIF/MT-3_S4 model). Although the theoretical and experimental Raman spectra exhibited different overall shapes, owing to computational limitations, it is clear that the peak near 511 cm^−1^ was markedly more intense for the apo-GIF/MT-3_S4 model than the apo-GIF/MT-3_S2 model. The normal mode vectors were evaluated, and the resulting assignments of these peaks are summarized in Table S1. The peaks mainly corresponded to the stretching and bending of disulfide and tetrasulfide bonds. The commensurate increase in peak intensity with the number of S–S bonds was consistent with the fact that the apo-GIF/MT-3_S4 model has several S–S and S–S–S–S bonds, while the apo-GIF/MT-3_S2 model has only S–S bonds. The Zn-binding models with/without persulfide also showed that the peaks around 500 cm^−1^ were almost lost in Zn_7_GIF/MT-3 without persulfide (Fig. S1 and Table S2), indicating that the peak near 488 cm^−1^ (Fig. 2F) for Zn_7_GIF/MT-3 corresponded to the S–S structure of persulfide. Taken together, these results suggest the existence of sulfane sulfur atoms in both Zn_7_GIF/MT-3 and apo-GIF/MT-3.

### Determination of sulfane sulfur atoms in Zn_7_GIF/MT-3

HPE-IAM is a relatively inert electrophile that allows the detection of sulfur atoms (e.g., H_2_S) by forming a bis-S-β-(4-hydroxyphenyl)ethyl acetamide (bis-S-HPE-AM) adduct (*8*). Our rationale was that if GIF/MT-3 is an SSBP, the interaction of HPE-IAM with Zn_7_GIF/MT-3 should eventually form a bis-S-HPE-AM adduct that can be quantified using LC-MS/MS with the stable isotope-labeled standard bis-S^34^-HPE-AM (Fig. 3A). Small molecules such as H_2_S were removed during the purification of Zn_7_GIF/MT-3 to exclude their contribution to the measured bis-S-HPE-AM adduct concentration. In a preliminary examination, a negligible amount of sulfane sulfur in Zn_7_GIF/MT-3 could be detected after 36 h incubation with HPE-IAM at 37°C. Stillman and coworkers reported that it was difficult for *N*-ethylmaleimide to access an apo-MT isoform at 37°C because of its folded structure, whereas heat treatment allowed such an electrophile to covalently bind the protein (*23*). Therefore, as expected, the amount of sulfane sulfur detected in Zn_7_GIF/MT-3 depended on the HPE-IAM concentration (plateauing at 5 mM), the amount of Zn_7_ GIF/MT-3 (up to 10 µM), and the reaction temperature and duration (Fig. 3B–D). We performed the reaction with 5 mM HPE-IAM at 60°C for 36 h. Under these optimized conditions, each MT isoform (each containing 20 cysteine residues) possessed approximately 20 sulfane sulfurs (Fig. 3E). None of the Cys-to-Ala mutants of GIF/MT-3 possessed sulfane sulfur (Fig. 3E), indicating that all 20 sulfane sulfurs were bound to cysteine residues of Zn_7_GIF/MT-3. Although the form of binding (e.g., 20 RSSH, 10 RSSSH, RSS_20_SR, 2 RSS_10_SR) was not identified, persulfide (20 RSSH) was suggested to be formed rather than polysulfide for reasons described in the discussion section.

**Fig. 3.**
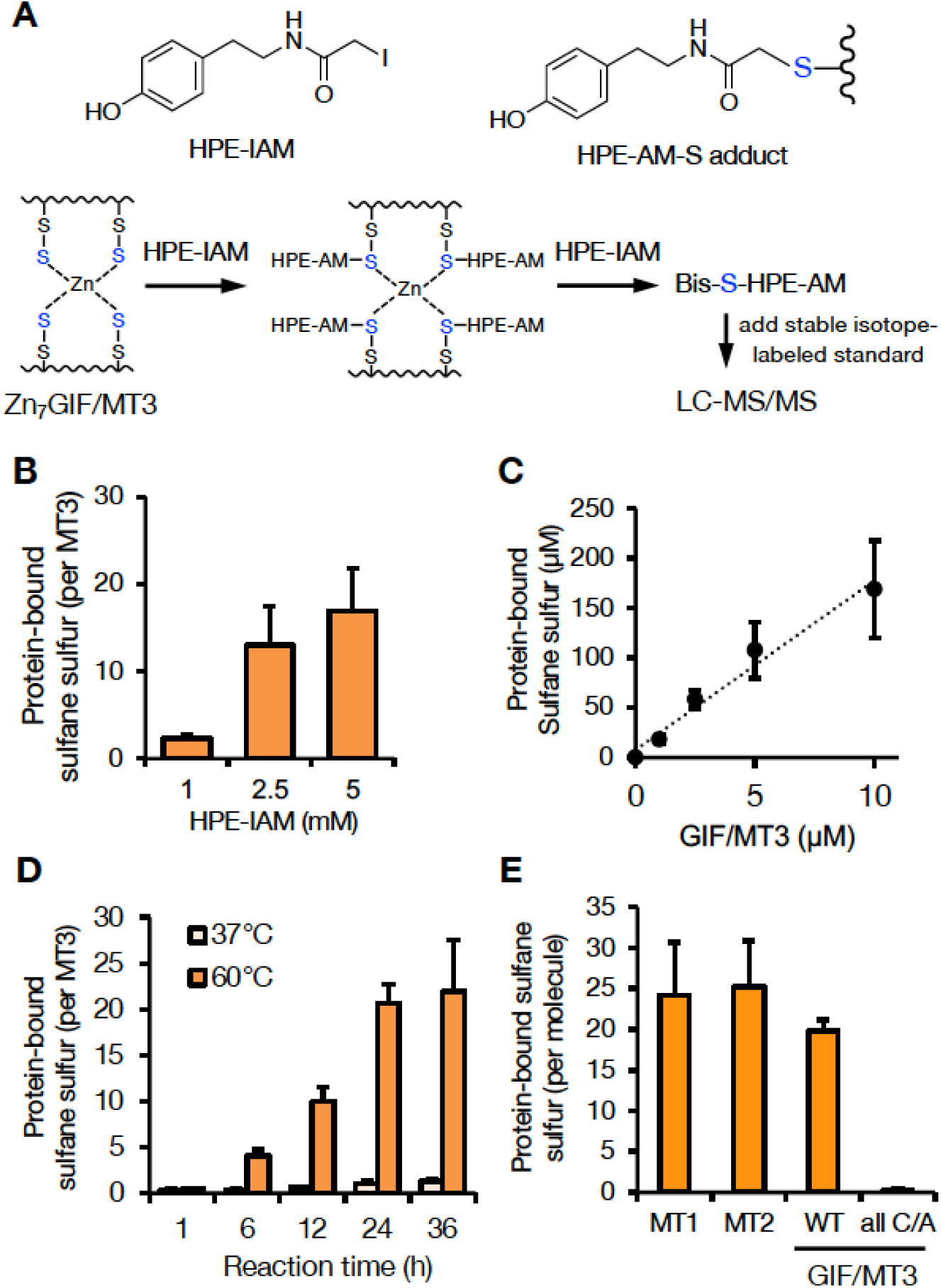
Sulfane sulfur assay optimization and quantification of MT sulfane sulfur content. (A) Schematic showing the detection of sulfane sulfur in Zn_7_GIF/MT-3. (B) Sulfane sulfur detected in Zn_7_GIF/MT-3 after incubation with the indicated concentrations of HPE-IAM at 60°C for 36 h. (C) Sulfane sulfur detected in Zn_7_GIF/MT-3 after incubation with 5 mM HPE-IAM at 60°C for 36 h. (D) Sulfane sulfur detected in Zn_7_GIF/MT-3 after incubation with 5 mM HPE-IAM at 37°C or 60°C for the indicated times. (E) Sulfane sulfur detected in human Zn_7_MT-1, Zn_7_MT-2, Zn_7_GIF/MT-3, wild-type (WT) Zn_7_GIF/MT-3, and apo-GIF/MT-3 with all Cys residues mutated to Ala (all C/A), each incubated with 5 mM HPE-IAM at 60°C for 36 h. Sulfane sulfur content was measured using LC-MS/MS. HPE-IAM, β-(4-hydroxyphenyl)ethyl iodoacetamide.

Because this assay was performed at relatively high temperatures (60°C), we also examined the sulfane sulfur levels of several mutant proteins using chemically synthesized α- and β-domains of GIF/MT-3 to eliminate false-positive results. As shown in Fig. S2A, sulfane sulfur (less than 1 molecule per protein) was undetectable in chemically synthesized α- and β-domains of GIF/MT-3, whereas several molecules of sulfane sulfur per protein were detected in recombinant α- and β-domains exhibited (Fig. S2B, left panel). These findings indicated that the sulfane sulfur detected in our assay was derived from biological processes executed during the production of GIF/MT-3 protein. We further analyzed mutant proteins with β-Cys-to-Ala and α-Cys-to-Ala substitutions and found that their sulfane sulfur levels were comparable with those of the α- and β-domains of GIF/MT-3, respectively (Fig. S2B, left panel). Additionally, Ser-to-Ala mutation did not affect the sulfane sulfur levels of GIF/MT-3. The zinc content of each mutant protein was also determined under these conditions (Fig. S2B, right panel).

### Redox-based GIF/MT-3 recycling system during oxidative stress

To explore the functional role of sulfane sulfur in GIF/MT-3, we examined the stability of sulfane sulfur in the protein with or without bound zinc. Freshly prepared Zn_7_GIF/MT-3 and apo-GIF/MT-3 possessed almost the same amount of sulfane sulfur (Fig. 4A). Unexpectedly, sulfane sulfur content in Zn_7_GIF/MT-3 remained unchanged for up to 28 days in 20 mM Tris-HCl (pH 7.5) at 37°C (data not shown). In contrast, in apo-GIF/MT-3, sulfane sulfur content decreased markedly within 12 h of incubation, and the addition of zinc blocked any further decrease (Fig. 4A). This suggests that zinc ions are rapidly re-coordinated by the persulfide group in apo-GIF/MT-3, thereby stabilizing sulfane sulfur atoms. To examine the possibility of forming intramolecular cysteine tetrasulfide, which is stable and cannot react with iodoacetamide in the absence of a reducing agent (*24*), apo-GIF/MT-3 left in 20 mM Tris-HCl (pH 7.5) for 36 h (i.e., oxidized apo-GIF/MT-3) was incubated with the reducing agent tris(2-carboxyethyl)phosphine (TCEP). The presence of free SH/SSH groups in oxidized apo-GIF/MT-3, determined using 5,5′-ditiobis-(2-nitrobenzoic acid) (DTNB) (*25*), was negligible but increased following TCEP incubation, leading to a complete recovery of sulfane sulfur atoms (Fig. 4B). These observations led us to assume that the time-dependent disappearance of persulfide in apo-GIF/MT-3 (Fig. 4A) was not due to the oxidative degradation of sulfane sulfur but rapid closure of a ring that can be cleaved by TCEP. Moreover, our method, based on TCEP-mediated reduction of tetrasulfide and subsequent trapping of sulfane sulfur atoms by HPE-IAM, was validated using the model compounds *N*-acetylcysteine-tetrasulfide (Fig. 5A) and diallyltetrasulfide (Fig. 5B). In the absence of TCEP, minimal amounts of bis-S-HPE-AM adducts were detected in all the compounds examined. However, incubation with TCEP resulted in the stoichiometric detection of sulfane sulfur but not oxidized-*N*-acetylcysteine and diallyl disulfide, suggesting that sulfane sulfur was stably trapped by HPE-IAM during the 36 h TCEP incubation. A possible mechanism of reaction between tetrasulfide compounds and HPE-IAM is shown in Fig. 5C. Thus, we confirmed that sulfane sulfur atoms of cysteine tetrasulfide in apo-GIF/MT-3 could be preserved after zinc release and oxidation.

**Fig. 4.**
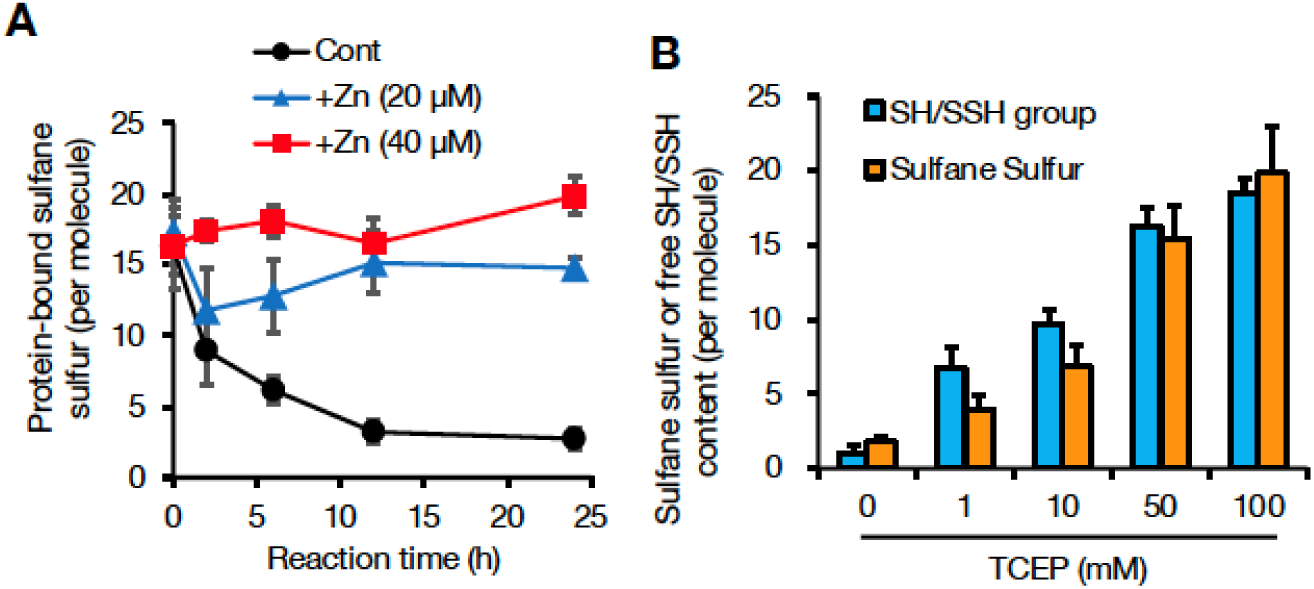
Sulfane sulfur stability in apo-GIF/MT-3 and its restoration by a reducing agent. (A) Stability of sulfane sulfur in apo-GIF/MT-3 incubated with or without (Cont) zinc. To prepare apo-GIF/MT-3, Zn_7_GIF/MT-3 was incubated in 0.1 M HCl for 30 min, then the buffer was replaced with 20 mM Tris-HCl (pH 7.5). To examine the stability of sulfane sulfur in apo-GIF/MT-3, freshly prepared apo-GIF/MT-3 (2 µM) with or without added zinc ions were incubated at 37°C for up to 24 h. (B) Effect of tris(2-carboxyethyl)phosphine (TCEP) on sulfane sulfur binding and free SH/SSH groups in oxidized apo-GIF/MT-3. To prepare oxidized apo-GIF/MT, Zn_7_GIF/MT-3 was incubated in HCl (0.1 N) at 37°C for 30 min and then replaced with 20 mM Tris-HCl (pH 7.5) buffer and incubated for 36 h at 37°C. The resulting oxidized apo-GIF/MT-3 protein (10 µM) was incubated with 0, 1, 10, 50, or 100 mM TCEP in 20 mM Tris-HCl (pH 7.5) at 37°C for 1 h, then low-molecular-weight molecules were removed by 3 kDa ultrafiltration for 6 times. Sulfane sulfur content was determined using LC-ESI-MS/MS, and the concentrations of free SH/SSH groups were measured using Ellman’s reagent.

**Fig. 5.**
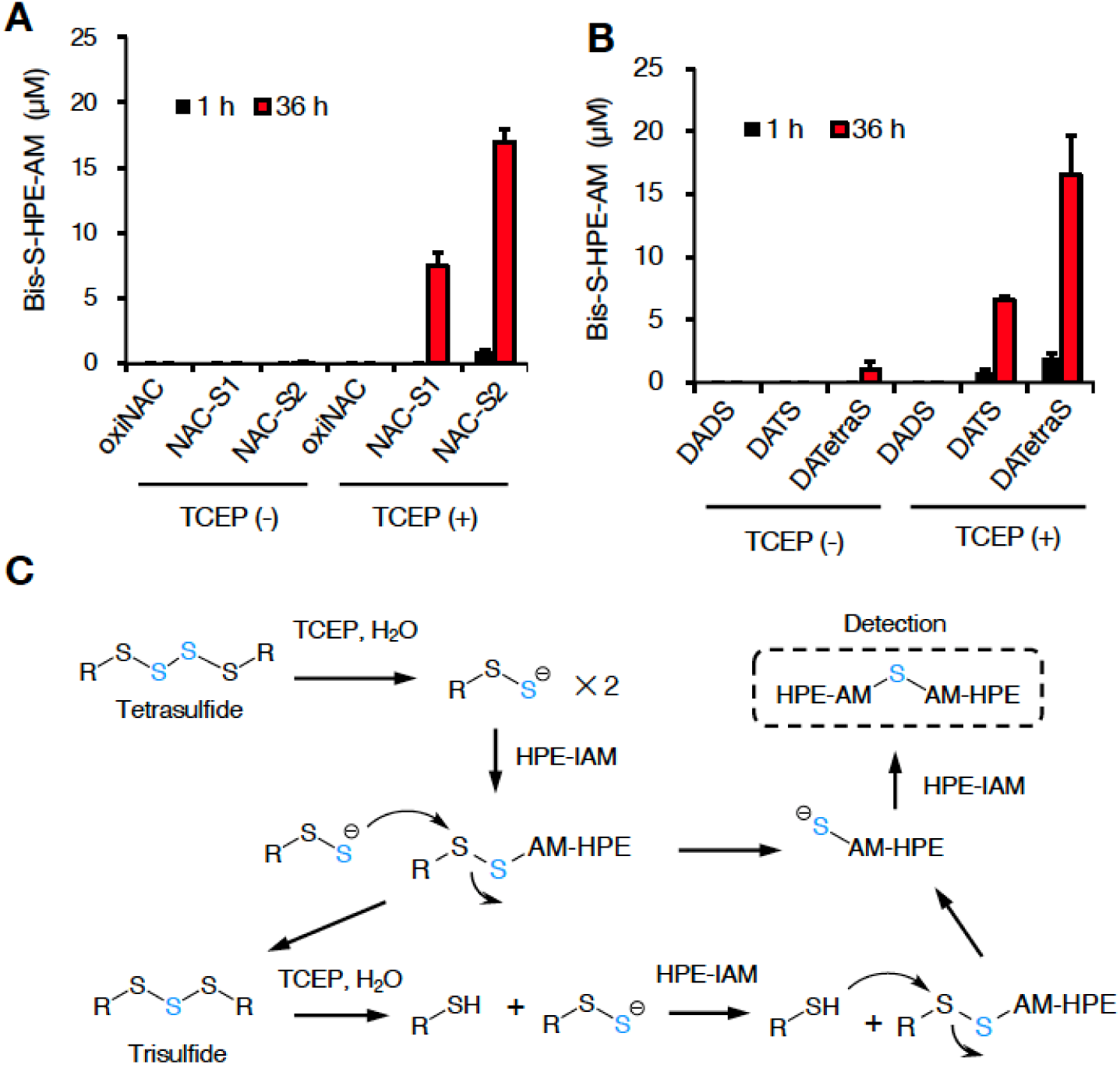
Reactivity of HPE-IAM with tetrasulfide derivatives as models of tetrasulfide bridges in apo-GIF/MT-3. (A) Reactivity of HPE-IAM with *N*-acetylcysteine (NAC) derivatives. Oxidized NAC (oxiNAC), NAC-trisulfide (NAC-S1), and NAC-tetrasulfide (NAC-S2) (each 10 µM) were incubated with HPE-IAM (5 mM) at 60°C for 1 or 36 h with or without TCEP (1 mM). (B) Reactivity of HPE-IAM with diallyl polysulfide derivatives. Diallyl disulfide (DADS), diallyl trisulfide (DATS), or diallyl tetrasulfide (DATetraS) (each 10 µM) were incubated with HPE-IAM (5 mM) at 60°C for 1 or 36 h with or without TCEP (1 mM). (C) Scheme showing possible reactions of tetrasulfide derivatives with HPE-IAM and TCEP. Bis-S-HPE-AM, bis-S-β-(4-hydroxyphenyl)ethyl acetamide.

Several reports have indicated that oxidative modification of MTs results in the release of zinc involved in zinc signaling (*26, 27*). Incubation with H_2_O_2_ and *S*-nitroso-*N*-acetylpenicillamine (SNAP), a nitrosonium ion donor, induced zinc release from Zn_7_GIF/MT-3 (Fig. 6A). Under these conditions, the numbers of free SH/SSH groups and sulfane sulfur atoms in GIF/MT-3 were also decreased by H_2_O_2_ and SNAP treatment. However, subsequent TCEP treatment nearly restored the original levels of free SH/SSH and sulfane sulfur (Fig. 6B). The persulfide in apo-GIF/MT-3 appeared to be resistant to TCEP-induced release of sulfane sulfur, although tetrasulfide was not. Overall, it seems likely that sulfane sulfur in Zn_7_GIF/MT-3 acts as a reserve of sulfur to be modified under physiological oxidative stress and a component of the redox-active closed-ring structure regulated by reductants (Fig. 6C).

**Fig. 6.**
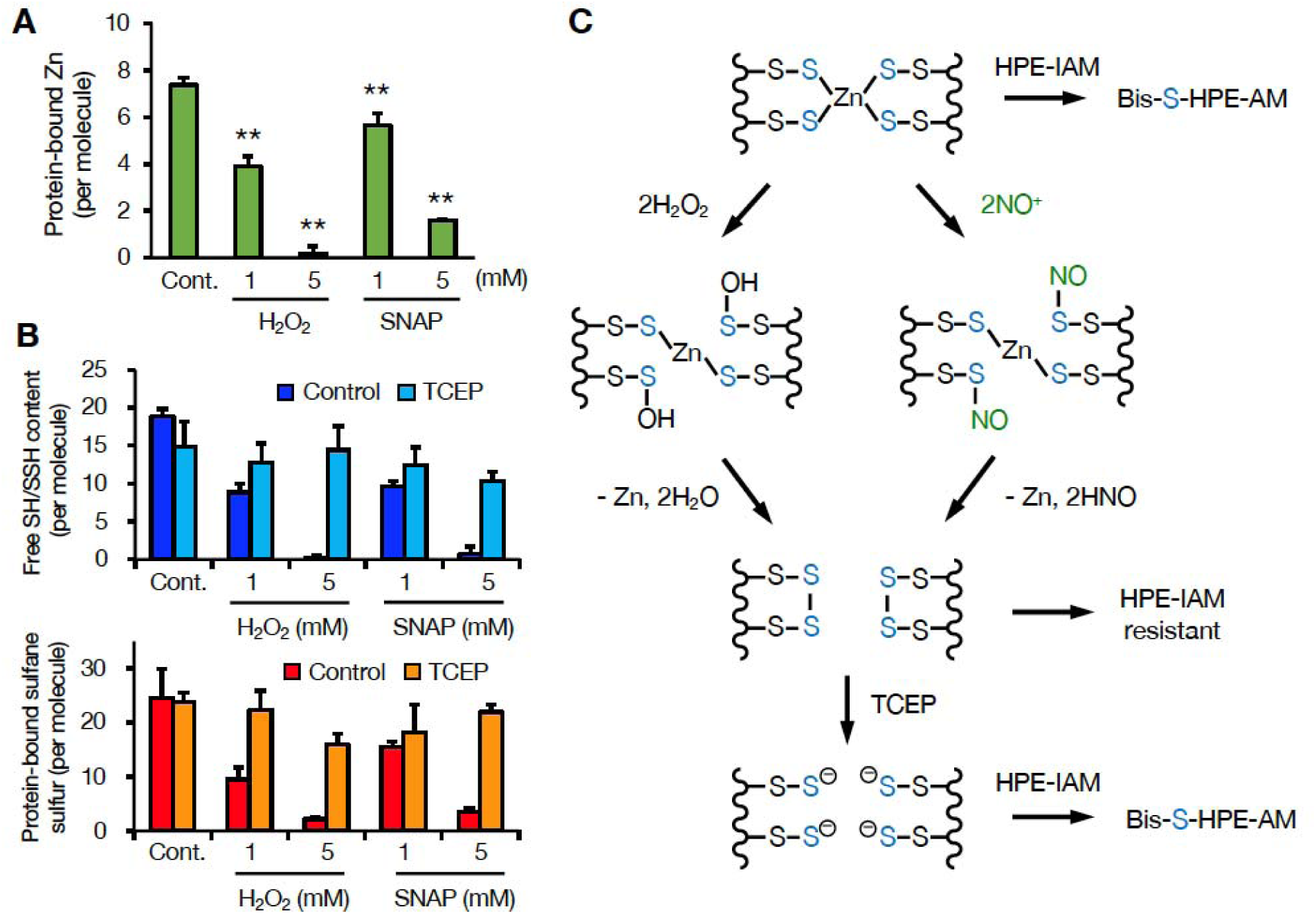
Redox-dependent release of zinc ions and recycling of sulfane sulfur in GIF/MT-3. (A) Quantitation of zinc ions released from Zn_7_GIF/MT-3 by H_2_O_2_ and *S*-nitroso-*N*-acetylpenicillamine (SNAP). To examine the release of zinc ions by H_2_O_2_ and SNAP, Zn_7_GIF/MT-3 (10 µM) was treated with H_2_O_2_ (1 or 5 mM) or SNAP (1 or 5 mM) in 100 mM Tris-HCl (pH 7.5) at 25°C for 30 min. After removing H_2_O_2_/SNAP using 3 kDa ultrafiltration for 4 times, free SH/SSH groups and sulfane sulfur content in GIF/MT-3 were determined. (B) Free SH/SSH content in Zn_7_GIF/MT-3, determined by H_2_O_2_ or SNAP treatment after incubation with TCEP. To examine the interaction of Zn_7_GIF/MT-3 with H_2_O_2_ or NO, Zn_7_GIF/MT-3 (10 µM) was incubated with H_2_O_2_ (1 or 5 mM) or SNAP (1 or 5 mM) in 100 mM Tris-HCl (pH 7.5) at 25°C for 30 min. After removing H_2_O_2_/SNAP using 3 kDa ultrafiltration for 4 times, the resulting proteins (5 µM) were incubated with TCEP (50 mM) in 100 mM Tris-HCl (pH 7.5) at 37°C for 1 h. After removing TCEP using 3 kDa ultrafiltration for 5 times, sulfane sulfur content was determined using LC-ESI-MS/MS and the concentrations of free SH/SSH groups were measured using Ellman’s reagent. (C) Proposed reactions between a zinc/persulfide cluster in GIF/MT-3 and H_2_O_2_ or NO.

The interaction of sulfane sulfur species with KCN to yield thiocyanate and thiol products (cyanolysis) has been used as evidence of the presence of protein hydropersulfides (*28*). Therefore, we used KCN to eliminate the sulfane sulfur atoms from Zn_7_GIF/MT-3. KCN treatment decreased the sulfane sulfur atom content of Zn_7_GIF/MT-3 by approximately 75% (Fig. 7A). After removing KCN, the reducing agent TCEP was subsequently added because intramolecular cysteine disulfide/tetrasulfide bridges can be formed under the condition. TCEP did not recover the level of sulfane sulfur in GIF/MT-3 (Fig. 7A), indicating that KCN indeed removed sulfane sulfur from GIF/MT-3. In addition to eliminating the sulfane sulfur atoms, KCN treatment also reduced the zinc content of GIF/MT-3 (Fig. 7B). To examine the role of sulfane sulfur in zinc retention, sulfane sulfur-diminished apo-GIF/MT-3 was incubated with zinc after TCEP treatment to reconstruct zinc-bound GIF/MT-3. Re-coordination of zinc ions to KCN-treated apo-GIF/MT-3 was incomplete compared with KCN-untreated apo-GIF/MT-3 (Fig. 7B), implying the contribution of sulfane sulfur to zinc binding in GIF/MT-3.

**Fig. 7.**
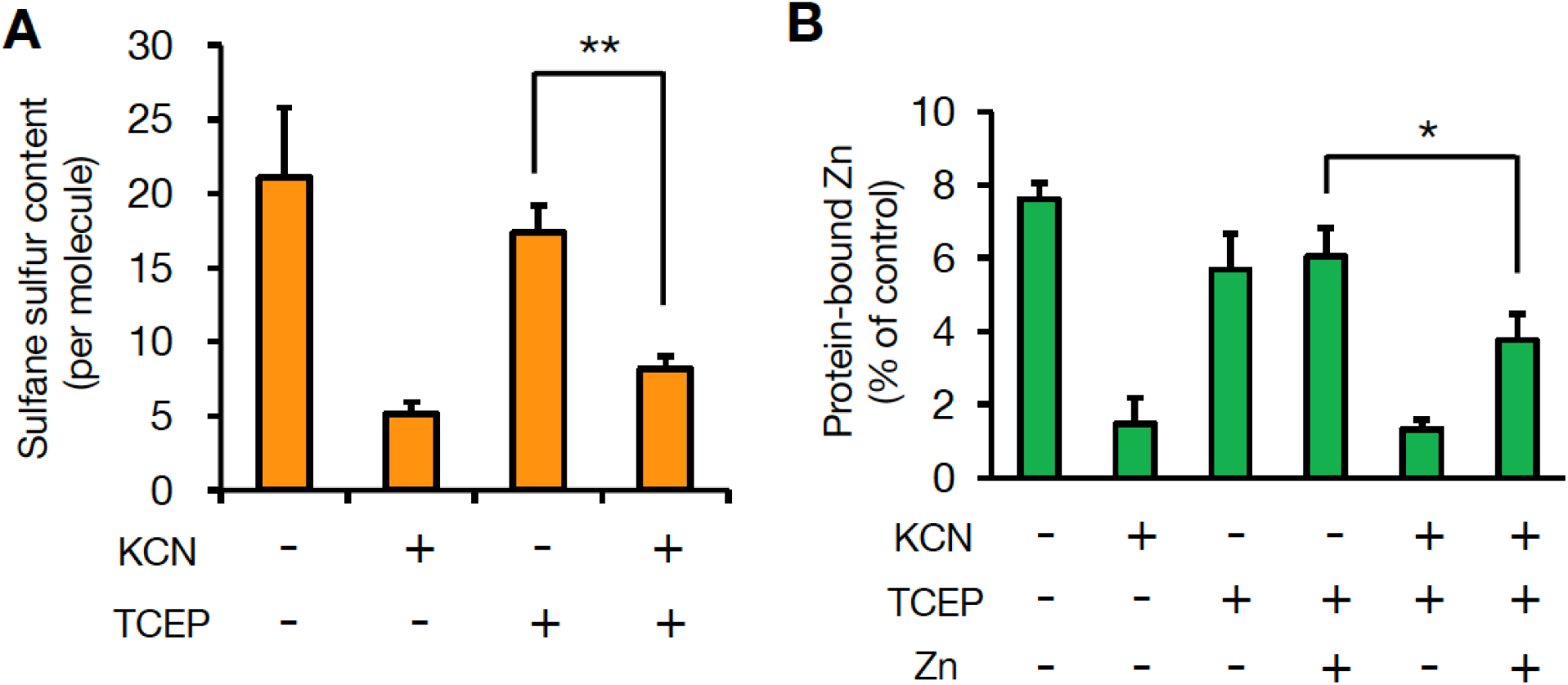
Contribution of sulfane sulfur in GIF/MT-3 to zinc binding. (A) To eliminate sulfane sulfur in Zn_7_GIF/MT-3 by cyanolysis, Zn_7_GIF/MT-3 (10 µM) was reacted with KCN (75 mM) in 100 mM Tris-HCl (pH 7.5) at 37°C for 14 h. After removal of KCN, the resulting protein was incubated with TCEP (10 mM) in 100 mM Tris-HCl (pH 7.5) at 37°C for 1 h. After removal of TCEP, sulfane sulfur content in GIF/MT-3 was determined using LC-ESI-MS/MS. (B) Comparison of zinc binding capacity of GIF/MT-3 before and after cyanolysis. Zn_7_GIF/MT-3 (10 µM) was incubated with KCN (75 mM) in 100 mM Tris-HCl (pH 7.5) at 37°C for 14 h. After removal of KCN, the resulting protein was incubated with TCEP (10 mM) in 100 mM Tris-HCl (pH 7.5) at 37°C for 1 h. After removal of TCEP, the resulting protein (5 µM) was incubated with zinc chloride (50 µM) in 50 mM Tris-HCl (pH 7.5) at 37°C for 1 h. Low-molecular-weight molecules were removed using 3 kDa ultrafiltration after each step. Protein-bound zinc content was determined using ICP-MS. \**p* < 0.05 and \*\**p* < 0.01.

### Reduction of apo-GIF/MT-3 by thioredoxin

Thioredoxin (Trx) is a master enzyme that reduces disulfide bonds in cellular proteins (*29*). Holmgren previously reported that *E*. *Coli* Trx predominantly catalyzes S–S bond cleavage of insulin (*K*_m_ ∼ µM) rather than low-molecular-weight substances such as cystine and oxidized GSH (*30*). Surprisingly, apo-GIF/MT-3 (*K*_m_ = 30 nM, *K*_cat_ = 31,536 min^−1^, *K*_cat_/*K*_m_ = 1051 × 10^6^ M^−1^min^−1^) was a much more efficient substrate than insulin (*K*_m_ = 1,192 nM, *K*_cat_ = 19,114 min^−1^, *K*_cat_/*K*_m_ = 16 × 10^6^ M^−1^min^−1^) (Fig. 8A and B), whereas Trx was unable to reduce Zn_7_GIF/MT-3 (Fig. 8C), as was the case for TCEP (Fig. 7A). Therefore, we hypothesized that the zinc/persulfide clusters in Zn_7_GIF/MT-3 may block the interaction of the protein with Trx and that the zinc ions bound to GIF/MT-3 may act as a repressor of Trx-mediated S–S bond cleavage. In addition, apo-GIF/MT-3 was a poor substrate for Trx-related proteins 14 (TRP14) and 32 (TRP32) (Fig. 8C). Furthermore, HPE-IAM trapping assay analysis confirmed that the Trx/Trx reductase (TR) system recovered sulfane sulfur content in apo-GIF/MT-3 (Fig. 8D) but not in the low-molecular-weight (<3 kDa) fraction (data not shown). These observations indicate that Trx is a powerful enzyme that cleaves the tetrasulfide bond in apo-GIF/MT-3 and that persulfide formation derived from apo-GIF/MT-3 seems to be resistant to Trx as well as TCEP. Notably, a small amount of sulfane sulfur in apo-GIF/MT-3 was observed even in the absence of Trx and TR, thereby supporting our conclusion (Fig. 2B–E) that some persulfides or their deprotonated forms exist in apo-GIF/MT-3. This may be a unique SSBP-related feature of GIF/MT-3.

**Fig. 8.**
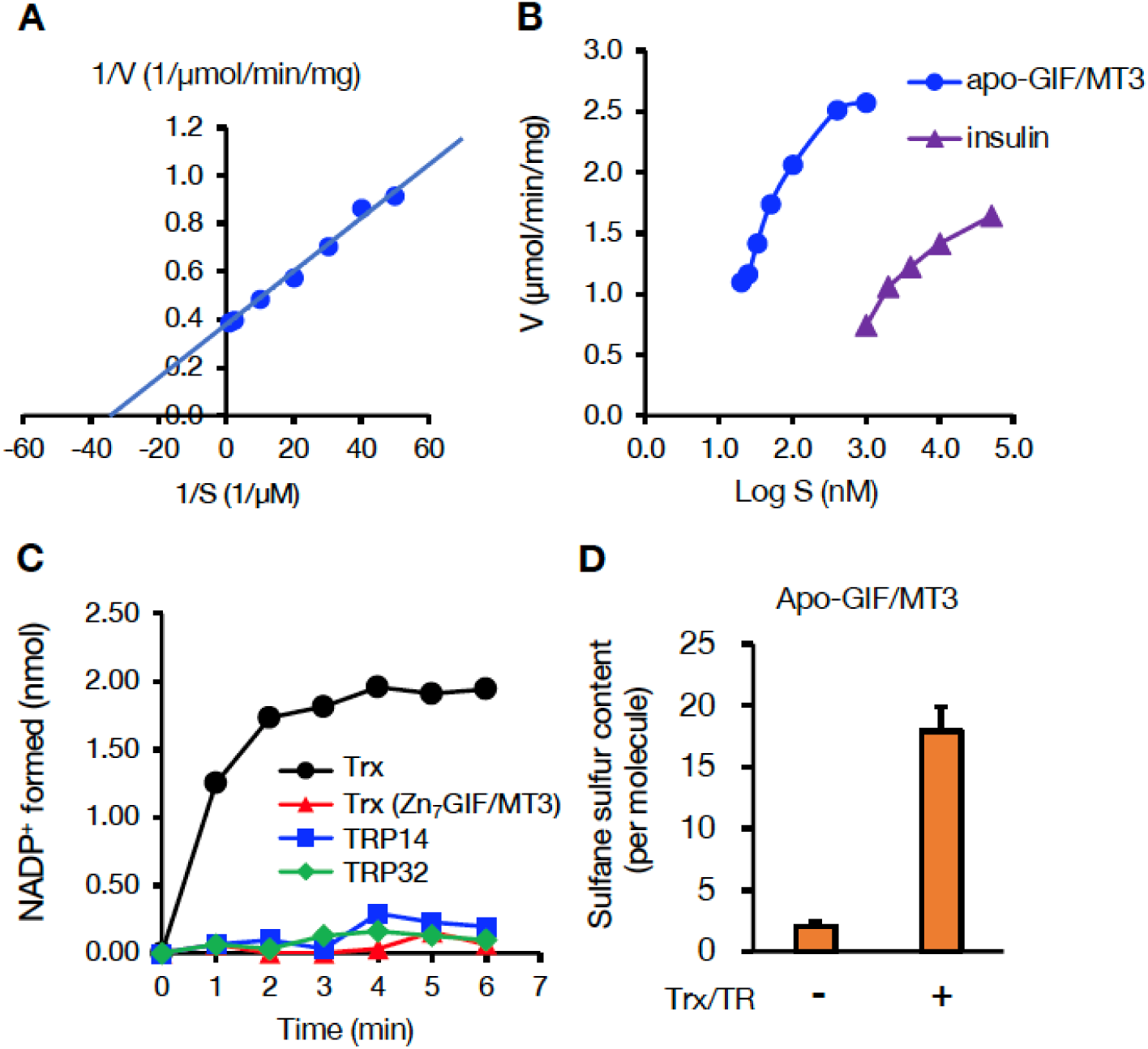
Reduction of apo-GIF/MT-3 by thioredoxin (Trx) and subsequent regeneration of sulfane sulfur. (A) Velocity (V) of Trx-catalyzed reduction of oxidized apo-GIF/MT-3 substrate (S). Oxidation of NADPH was followed by measuring the absorbance of NADPH at 340 nm. (B) Comparison of substrate reduction by NADPH and the thioredoxin system. (C) NADP^+^ formation upon incubation of: oxidized apo-GIF/MT-3 with Trx/TR, TRP14/TR, or TRP32/TR; and Zn_7_GIF/MT-3 with Trx/TR. (D) Regeneration of sulfane sulfur in oxidized apo-GIF/MT-3 after incubation with the Trx/TR system. TR, Trx reductase; TRP14, Trx-related protein 14; TRP32, Trx-related protein 32.

### 3D modeling of GIF/MT-3 with sulfane sulfur atoms

We generated a 3D homology model of human Zn_7_GIF/MT-3 using the Molecular Operating Environment (MOE) software and the Protein Data Bank (PDB) structures 4MT2 and 2F5H as templates (Fig. S3). Then we created a 3D structure of sulfane sulfur-bound Zn_7_GIF/MT-3 (one sulfane sulfur per cysteine residue), Zn_7_S_20_GIF/MT-3 (Fig. 9A), which was also used for Raman spectra modeling (Fig. 2B-E). The predicted 3D structure of Zn_7_S_20_GIF/MT-3 was almost the same as that of Zn_7_GIF/MT-3 (Fig. 9A), with the root-mean-square deviation of atom positions being only 0.789 Å. Similar results were obtained for Zn_7_MT1 and Zn_7_MT2 (Fig. S4). Zn_7_GIF/MT-3 contains a cyclohexane-like Zn_3_Cys_9_ cluster in the β domain and a bicyclononane-like Zn_4_Cys_11_ cluster in the α domain (*31*). The structures of both clusters were mostly maintained even when all thiol groups were changed to persulfides by adding one sulfane sulfur atom to each (Fig. 9B). Schematic structures of the generated Zn_7_GIF/MT-3 with or without sulfane sulfurs are shown in Fig. S5. Addition of two sulfane sulfur atoms, corresponding to cysteine trisulfide, disrupted each cluster structure (cf. Fig. S6 and Fig. 9B). When they contained one sulfane sulfur in each cysteine, the thermostability scores of MT1, MT2, and GIF/MT-3 in the presence of zinc ions were higher than those in the absence of zinc ions (Table S3), indicating that these ions are key to the thermostability of MT isoforms, including those containing sulfane sulfur atoms. The thermostabilities and Cd_7_, Cu_7_, Hg_7_, and Zn_7_ binding affinities of S_20_GIF/MT-3 were more favorable than the corresponding values of the sulfane sulfur-free protein (Table 1). Conversely, placing more than one sulfane sulfur on each cysteine residue decreased the thermostability and zinc binding affinity (Fig. 9C). Collectively, these results indicate that zinc ions contribute to protection against persulfide oxidation and MT thermostability, while sulfane sulfur atoms participate in cysteine tetrasulfide formation, enhancement of metal binding affinity. Therefore, our study provides evidence for an interdependence of zinc and sulfane sulfur and for unique structural and functional roles of the persulfide groups in GIF/MT-3.

**Fig. 9.**
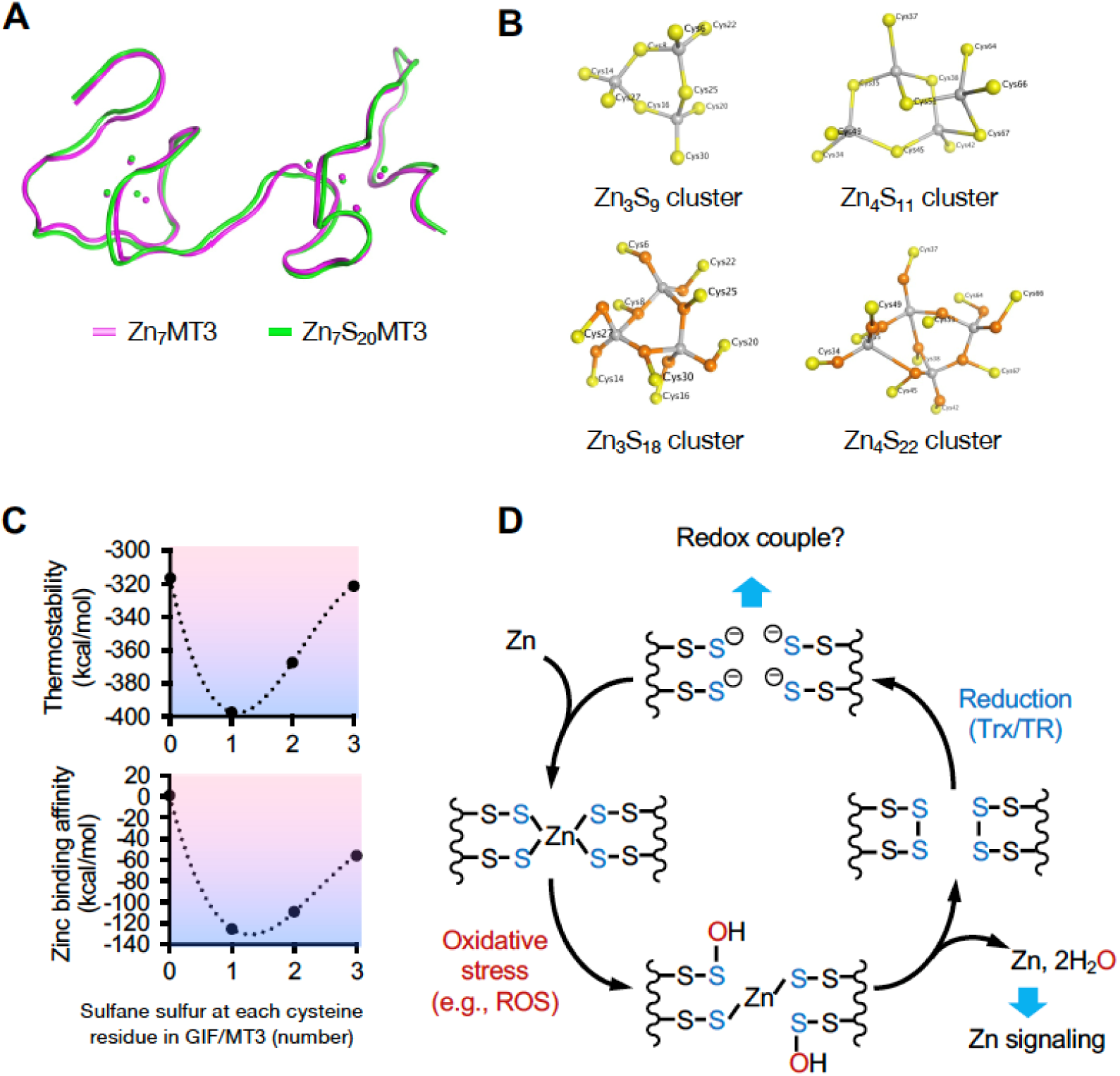
Structural modeling of sulfane sulfur in GIF/MT-3 using MOE, and a reaction scheme for sulfane sulfur-based zinc/persulfide cluster. (A) Comparison of three-dimensional structures of Zn_7_GIF/MT-3 (pink) and Zn_7_S_20_GIF/MT-3 (green). (B) Cyclohexane-like Zn_3_Cys_9_ cluster in the GIF/MT-3 homology model, and bicyclononane-like Zn_4_Cys_11_ cluster derived from PDB structure 2F5H with (lower) or without (upper) sulfane sulfur. Yellow, orange, and gray spheres indicate cysteine residues, sulfane sulfur, and zinc ions, respectively. (C) Thermostability and zinc binding affinity scores of GIF/MT-3 with different numbers of sulfane sulfurs at each cysteine residue. (D) A proposed model for redox-dependent hold-and-release regulation of zinc ions by GIF/MT-3.

**Table 1.** Thermostability and metal binding affinity scores of growth inhibitory factor (GIF)/metallothionein-3 (MT-3) with or without sulfane sulfur. Values were calculated using the Protein Design module of the Molecular Operating Environment (MOE) software.

| Metal | Sulfane sulfur | Affinity (kcal/mol) | Stability (kcal/mol) |
| --- | --- | --- | --- |
| Zn<sub>7</sub> | 0 | 11 | -302 |
|  | 20 | -154 | -407 |
| Cd<sub>7</sub> | 0 | 12 | -282 |
|  | 20 | -82 | -344 |
| Hg<sub>7</sub> | 0 | 13 | -276 |
|  | 20 | -112 | -354 |

To confirm the presence of SSBPs in mouse brain, we attempted to isolate them from the high-molecular-weight fraction (>3 kDa) of the cytosol using diethylaminoethyl (DEAE)-Sepharose CL-6B column chromatography. Surprisingly, an abundance of SSBPs were detected, and approximately half of them were tightly bound to the column and eluted in buffer containing 0–0.4 M NaCl (Fig. 10A). Although SSBPs from other proteins with iron/sulfur clusters may have also been detected, this possibility remains to be explored in future studies. Eluate corresponding to the peak concentration of SSBPs (fractions 40 to 44) was further separated using Sephacryl S-100 column chromatography (Fig. 10B), which resolved two major SSBP-related peaks. We collected eluate corresponding to the latter (fractions 40 to 43), which contained proteins with high SSBP content and a mass of approximately 13 kDa, even though its total protein concentration was low. These proteins were then separated using Blue Sepharose chromatography. While proteins that bound tightly to the Blue Sepharose resin did not contain sulfane sulfur, an SSBP that eluted in fractions 3 to 5 migrated as a single band (16 kDa) using SDS-PAGE (data not shown). This 16 kDa SSBP was confirmed to be GIF/MT-3 using nano ultra-performance LC-MS/MS (Table S4).

**Fig. 10.**
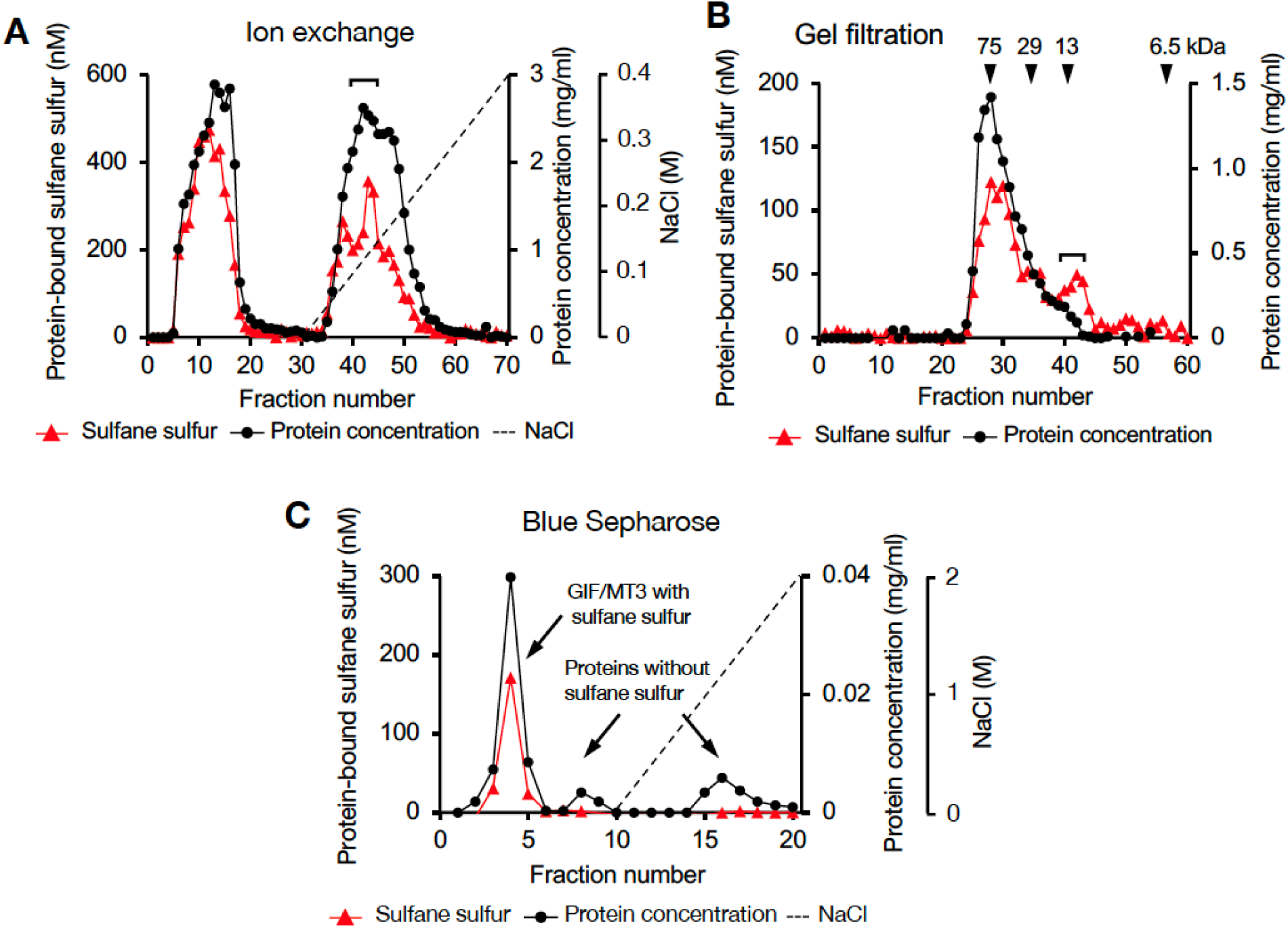
Separation of sulfane sulfur-binding proteins from mouse brain cytosol using column chromatography. (A) Diethylaminoethyl Sepharose CL-6B column. (B) Sephacryl S-100 column. (C) Blue Sepharose column. Triangles, closed circles, and dotted lines indicate sulfane sulfur, protein, and NaCl concentrations, respectively. Portions of each fraction were incubated with 5 mM of HPE-IAM in 20 mM Tris (pH 7.5) at 37°C for 30 min and the sulfane sulfur content was determined from the bis-S-HPE-AM adduct concentration measured using LC-MS/MS. Protein concentration was determined using the bicinchoninic acid assay. Isolation of sulfane sulfur-binding protein was performed as described in the Experimental procedures.

## Discussion

MT isoforms are believed to possess a zinc/thiolate cluster characterized by strong Zn–S bonds but facile release of zinc ions occurs under oxidative stress (*12, 13*). Here, we developed a reliable assay involving prolonged (36 h) incubation with HPE-IAM at 60°C to extract sulfane sulfur from Zn_7_GIF/MT-3 and isotope-dilution LC-MS/MS analysis (Fig. 3). The present study provided evidence that MTs are SSBPs containing a sulfane sulfur atom on each of their 20 cysteine residues, which form a zinc/persulfide cluster. Although Capdevila et al. previously showed evidence for existence of sulfide ions in recombinant MT1, MT2, and MT4, they detected only one to four sulfide ions liberated from MTs as H_2_S gas using gas chromatography–flame photometric detection in strongly acidic conditions (*15*). These observations suggest that unlike our assay, this method is likely to underestimate the sulfane sulfur content of proteins. While our assay has difficulty detecting sulfane sulfur in oxidized polysulfide bridges such as cysteine tetrasulfide, which is supported by a recent observation (*24*), a TCEP cleavage step enabled this problem to be overcome (Fig. 5C). However, FT-ICR-MALDI-TOF/MS analysis failed to detect sulfur modifications in GIF/MT-3 (Fig. 1B), suggesting that sulfur modifications in the protein were dissociated during laser desorption/ionization. Therefore, we postulate that the small amount of sulfur detected in oxidized apo-GIF/MT-3 is derived from the effect of laser desorption/ionization rather than any actual modification of the minority component.

Sulfane sulfur modification of MTs appears to be universal because similar sulfane sulfur contents were observed in recombinant human MT-1, MT-2, and GIF/MT-3 (Fig. 3E). Although MTs have been mainly studied in vertebrates, their diversity and distribution have been widely reported (*32*). So far, three functional MT forms (reduced apoprotein, oxidized apoprotein, and metalated protein) are known. In this current study, we propose that sulfane sulfur is another key factor involved in regulation of MT function that may have major implications for several biological functions.

Oxidation of sulfane sulfur in GIF/MT-3 is a fascinating mechanism of zinc release that does not involve direct thiol oxidation. There is little doubt that zinc release from the zinc/persulfide cluster in GIF/MT-3 will be sensitive to mild oxidative stress because the p*K*_a_ value of cysteine persulfide is lower than that of cysteine (*33*). However, formation of cysteine tetrasulfides following zinc release from the oxidized GIF/MT-3 (Fig. 9D) represents a paradigm shift in GIF/MT-3 biochemistry. The sulfane sulfur atoms of MT-tetrasulfide are stably retained and can reacquire zinc after they are reduced, as shown in Fig. 9D. Tetrasulfides, whose presence in apo-GIF/MT-3 was shown (Fig. 1C and Fig. 2A), are presumably formed in response to structural frustration of the disulfide form, as reported for the sulfide-responsive transcriptional repressor, SqrR (*24*). In our preliminary study, we failed to directly identify cysteine tetrasulfide during reaction of apo-GIF/MT-3 with pronase because alkaline hydrolysis of polysulfide makes it unstable in water (*34, 35*). We therefore speculate that cysteine tetrasulfide groups exist in an acidic local environment of apo-GIF/MT-3 and that Trx rapidly interacts with and cleaves tetrasulfide S–S bonds because of their markedly low p*K*_a_ values, resulting in the regeneration of persulfide groups.

It is believed that the function of the sulfane sulfur moiety in proteins is to protect cysteine thiols from irreversible oxidation (e.g., to sulfinic acid and sulfonic acid) because such oxidative modifications of cysteine persulfides can be reversed by Trx-mediated reduction (*36*). In other words, sulfane sulfurs act as “sacrificial” sulfur atoms. However, in this study, we discovered that sulfane sulfur has a higher affinity for zinc than thiol (Fig. 9C and Table 1) and is stored in non-sacrificial tetrasulfide groups when oxidized (Fig. 9D). As a result, tetrasulfides in oxidized-apo-GIF/MT-3 are reduced by the Trx system, with Trx being regenerated to a reduced form by NADPH and TR, thereby enabling reacquisition of zinc ions by the persulfide moiety. This suggests that the persulfide moiety in GIF/MT-3 appears to be relatively stable against Trx reduction. In contrast, Trx has been proposed to reduce the persulfide moiety of PTP1B (*37*) and albumin (*38, 39*). A possible explanation for this discrepancy is that apo-GIF/MT-3-persulfide is rapidly changed into a different conformation that is topologically resistant to Trx reduction. In other words, Trx may exhibit substrate specificity.

While our proposed mechanism of MT redox regulation is consistent with that proposed by Maret et al. (*13*), discovery of sulfane sulfur in MTs explains unknown features of MT redox biochemistry. Maret and coworkers showed that approximately 50% of MT existed in the apo form in the rat brain, although they did not examine specific isoforms (*40*). A reasonable explanation for this observation is that reduced apo-MT can undergo redox-coupled reactions with oxidized proteins (*41*), leading to the formation of apo-MT-tetrasulfide, which loses its zinc ion-binding capability. GIF/MT-3 is a constitutive form predominantly expressed in the brain (*42*) and protects against Alzheimer’s disease (*43, 44*). Notably, several studies have indicated that Trx acts as a protection factor in Alzheimer’s disease (*45, 46*). Although the involvement of GIF/MT-3 and Trx in regulating this disorder has not been elucidated, Trx-mediated reduction of apo-GIF/MT-3 may lead to the reduction of unknown proteins that might suppress the onset of Alzheimer’s disease because of the high antioxidant capabilities of apo-MT persulfides. Further studies are required to identify the redox-coupled protein dynamics associated with the reduced form of apo-GIF/MT-3 and their relevance to the molecular basis of Alzheimer’s disease.

To our knowledge, this is the first study to perform 3D structural modeling of an SSBP with or without sulfane sulfur. Our structural modeling studies revealed that, like Zn_7_GIF/MT-3, the seven zinc ions of Zn_7_S_20_GIF/MT-3 were tetrahedrally coordinated by the array of 20 sulfane sulfurs but not by cysteine thiols, thereby forming two zinc/persulfide clusters (Fig. 9B). As shown in Figs. 9A and S3, cyclohexane-like Zn_3_S_9_ and bicyclononane-like Zn_4_S_11_ clusters were conserved even in the presence of 20 sulfane sulfurs without affecting the overall structure of the MTs. Nevertheless, sulfane sulfur is important for both metal binding affinity and protein stability (Table 1) and thus MT function. We suggest that the 20 sulfane sulfurs are evenly distributed among 20 cysteine residues in GIF/MT-3 for the following reasons: i) zinc binding of Zn_7_GIF/MT-3 was suggested to involve a C–S–S–Zn rather than Zn–S–Zn structure (Fig. 2A); ii) addition of two or three sulfane sulfurs to each cysteine residue in Zn_7_GIF/MT-3 decreased its stability (Fig. 9C); iii) the polysulfide form of sulfane sulfur seems to have difficulty maintaining zinc/sulfur clusters in GIF/MT-3 (Fig. S6). However, one of the limitations of our study is that we did not directly obverse such zinc-persulfide cluster itself.

Because protein sulfuration occurs during nascent protein translation, SSBPs appear to be ubiquitous in cells. In fact, we previously reported that a variety of cellular proteins were per/poly-sulfidated, as determined using a tag-switch-tag assay. We have recently characterized several SSBPs such as GSH S-transferase P1 (GSTP1) (*9*), dynamin-related protein 1 (Drp1), alcohol dehydrogenase 5 (ADH5), glyceraldehyde-3-phosphate dehydrogenase (GAPDH), ethylmalonic encephalopathy 1 (ETHE1) (*8*), and now MT1, MT2, and GIF/MT-3. As shown in Fig. 10, the present assay also confirmed that there are numerous SSBPs in addition to GIF/MT-3 in the cytosol of the mouse brain. Notably, 3,207 zinc-binding proteins in the human proteome have been identified with three and four zinc ligands, with 40% of the latter category consisting of Cys4 coordination (*47*). Once oxidants react with thiols in a zinc finger domain, zinc is released from the coordination site, resulting in inhibition of the zinc finger protein function. However, disulfide formation in the zinc finger domain is reversed by reducing agents such as GSH. We speculate that sulfane sulfur is a key component of such a “zinc redox switch”, enabling it to provide high-affinity for zinc and protection against excess oxidative/electrophilic stress. Supporting our notion, zinc finger proteins such as tristetraprolin (*48*), androgen receptor (*49*), and prolyl hydroxylase (*50*) were reported to undergo persulfidation by H_2_S. In this context, H_2_S inhibits enzymatic activity of tristetraprolin and androgen receptor, but increases enzymatic activity of prolyl hydroxylase. A possible explanation for this difference is that excess exogenous H_2_S may cause protein polysulfidation, thereby disturbing the native protein structure. Our findings provide structural and mechanistic insights into the role of sulfane sulfur in hold-and-release regulation of zinc ions by zinc-binding proteins. A future research goal is to investigate the universal role of sulfane sulfur atoms in redox regulation of all zinc-finger proteins.

In conclusion, we have provided evidence that S-sulfuration based on addition of sulfane sulfur plays a central role in hold-and-release regulation of zinc by GIF/MT-3. Our study has further revealed a fascinating redox-dependent switching mechanism of a zinc/persulfide cluster involving formation of a cystine tetrasulfide bridge. The biological significance of sulfane sulfur in MTs lies in its ability to 1) contribute to metal binding affinity, 2) provide a sensing mechanism against oxidative stress, and 3) aid in the regeneration of the protein. We believe that our findings open new directions of research in redox and metals biology.

## Materials and Methods

### Materials

DEAE Sepharose CL-6B, Sephacryl S-100, Blue Sepharose 6 Fast Flow, Glutathione Sepharose 4 Fast Flow, and Benzamidine Sepharose 4 Fast Flow were obtained from GE Healthcare (Uppsala, Sweden). HPE-IAM was obtained from Molecular Biosciences (Boulder, CO, USA). Human recombinant thioredoxin and insulin were purchased from Wako Pure Chemical Industries (Osaka, Japan). TR from rat liver (TrxB) was obtained from Sigma-Aldrich (St. Louis, MO, USA). pGEX-4T-1 cells were obtained from GE Healthcare. Chemical synthesis of α-domain and β-domain of GIF/MT-3 proteins was carried out by HiPep Laboratories (Kyoto, Japan) using conventional solid-phase synthesis, followed by purification and characterization using LC–MS. The amino acid sequences of the proteins were identical to those shown in Fig. S7. All other reagents and chemicals used were of the highest grade available.

### Protein expression and purification

Vector pEX-K4J2 containing human cDNA of MT-1A, MT-2, GIF/MT-3, or GIF/MT-3 mutants (all Cys-to-Ala, α-domain, β-domain, βCys-to-Ala, αCys-to-Ala, or all Ser-to-Ala) between the *Bam*HI and *Xho*I sites was obtained from Eurofins Genomics (Tokyo, Japan). The cDNAs were excised using *Bam*HI and *Xho*I and ligated into the corresponding sites of pGEX-4T-1, a glutathione *S*-transferase (fusion expression vector. An overnight culture of *E*. *Coli* BL21 containing 1% v/v pGEX-4T1-1/cDNA vector in fresh Luria-Bertani medium was grown at 37°C for 2 h, then ZnCl_2_ (final concentration of 500 µM) and isopropyl-1-thio-β-D-galactopyranoside (final concentration of 100 µM) were added to induce the expression of the fusion protein. After 5 h incubation at 37°C, cells were pelleted by centrifugation at 5,000 × *g* for 10 min at 4°C and resuspended in 5% of the original volume of buffer A (20 mM Tris-HCl [pH 7.5], 150 mM NaCl), then lysed using mild sonication at 4°C. Triton X-100 was added to a final concentration of 1% v/v, and the suspension was mixed gently at approximately 20°C for 1 h to facilitate protein solubilization. After centrifugation at 105,000 × *g* for 1 h, the supernatant was applied at a flow rate of 2 mL/min to a Glutathione Sepharose column (5.5 cm × 1.5 cm i.d.) pre-equilibrated with buffer A. The column was washed with 100 mL of buffer A, then syringe-filled with 10 mL thrombin solution (400 U/mL in buffer A; Wako, Osaka, Japan) and incubated overnight at room temperature. After incubation, the target proteins and thrombin were eluted using buffer A. The eluate was filtered using a 3 kDa Amicon Ultra centrifugal filter unit (Millipore) following centrifugation with buffer C (20 mM Tris-HCl [pH 7.5], 500 mM NaCl) to concentrate the fractions and exchange the buffer. To remove thrombin, the concentrated sample was applied at a flow rate of 2 mL/min to a Benzamidine Sepharose 4 Fast Flow column (3.9 cm × 1.5 cm i.d.) pre-equilibrated with buffer C. The eluted sample was filtered eight times using a 3 kDa Amicon Ultra centrifugal filter unit with buffer B to exchange the buffer and remove small molecules, then stored at −80°C. The amino acid sequences of the obtained recombinant proteins are shown in Fig. S7.

### Protein assay

The cytoplasmic protein concentration in mouse brain was determined using the bicinchoninic acid assay with bovine serum albumin as a standard. The concentrations of GIF/MT-3 protein were determined by measuring the absorbance of apo-GIF/MT-3 at 220 nm using an extinction coefficient of 53,000 M^−1^cm^−1^ (*51*). To produce apo-GIF/MT-3, Zn_7_GIF/MT-3-containing buffer was exchanged with 0.1 M HCl by ultrafiltration. Briefly, protein solution (0.5 mL) was added to an Amicon Ultra centrifugal filter unit. After 30 min centrifugation at 14,000 × *g*, the filtrate was discarded and 0.1 M HCl (0.45 mL) was added to the retentate. This procedure was repeated two times, and the final retentate was exchanged with 20 mM Tris-HCl (pH 7.5) buffer and used for further studies.

### Protein isolation

Animal housing, husbandry, and euthanasia were conducted according to the guidelines of the Animal Care and Use Committee of the University of Tsukuba. Approximately 50 C57BL/6 mice (10–20 weeks-old, ≈1:1 male/female), kindly provided by Prof. S. Takahashi (University of Tsukuba), were anesthetized by intraperitoneal injection of 200 mg/kg phenobarbital. Their brains were perfused with cold saline and then homogenized with four volumes of buffer A (20 mM Tris [pH 7.5], 150 mM NaCl). The homogenate was centrifuged at 9000 × *g* for 10 min at 4°C and the resulting supernatant was centrifuged at 105,000 × *g* for 1 h to obtain the cytosol. The cytosolic fraction was filtered and buffer-exchanged 27 times using a 3 kDa Amicon Ultra centrifugal filter unit and centrifugation at 5,000 × *g* for 30 min at 4°C with buffer B (20 mM Tris [pH 7.5]). To isolate SSBPs, the resulting high-molecular-weight fraction, containing 442 mg protein in 69 mL solution, was applied to a DEAE Sepharose CL-6B column (4.1 cm × 2.5 cm i.d.) pre-equilibrated with buffer B. The column was washed with buffer B at a flow rate of 1 mL/min, then SSBPs were eluted using 200 mL buffer B with a linear gradient of 0–0.4 M NaCl and 5 mL fractions were collected. The major SSBP-containing fractions (fractions 40–44 in Fig. 10A) were filtered using a 3 kDa Amicon Ultra centrifugal filter unit and centrifugation at 14,000 × *g* for 30 min at 4°C with buffer A seven times to concentrate the protein and exchange the buffer. The concentrated fraction (4.5 mL) was applied to a Sephacryl S-100 column (71 cm × 2.5 cm i.d.) previously equilibrated with buffer A and eluted with buffer A at a flow rate of 1 mL/min and 5 mL fractions were collected The major SSBP-containing fractions with low total protein concentration (fractions 40–43 in Fig. 10B) were combined and filtered, as described above, with buffer B. The concentrated fraction (1 mL) was applied to a Blue Sepharose column (8.4 cm × 2.5 cm i.d.) previously equilibrated with buffer B. The loaded column was washed with buffer B, then SSBPs were eluted a flow rate of 1 mL/min using 200 mL buffer B with a linear gradient of 0–0.4 M NaCl and collected in 10 mL fractions. Eluate containing SSBP (fractions 3–5 in Fig. 10C) was concentrated to a volume of 0.2 mL using Amicon Ultra centrifugal filter units. The resulting material was used for sulfane sulfur detection, SDS-PAGE with silver staining, and western blotting. All operations were performed at 4°C. Protein-bound sulfane sulfur was measured by determining bis-S-HPE-AM concentration after incubation of the protein with HPE-IAM. The incubation mixture (100 µL) consisted of 5 mM HPE-IAM and protein in buffer B. The reaction was performed at 37°C for 30 min and the yield of the bis-S-HPE-AM adduct was determined using liquid chromatography-electrospray ionization-tandem mass spectrometry (LC-ESI-MS/MS), as described later in detail.

### Identification of proteins

To identify SSBP in the brain, the isolated protein (75 µg) was incubated with 20 mM dithiothreitol in 50 mM ammonium bicarbonate at 50°C for 1 h, then incubated with 50 mM iodoacetamide in 50 mM ammonium bicarbonate at room temperature for 30 min, and finally digested with trypsin (1.2 µg) at 37°C overnight. The tryptic digests (2.5 µL) were loaded in direct injection mode onto a nanoAcquity ultra-performance LC system (Waters, Milford, MA, USA) equipped with a BEH130 nanoAcquity C_18_ column (100 mm × 75 μm, 1.7 μm i.d.) held at 35°C. Mobile phases A (0.1% v/v formic acid) and B (0.1% v/v formic acid in acetonitrile) were linearly mixed at a flow rate of 0.3 μL/min using a gradient system as follows: 3% B for 1 min; linear increase to 40% B over 65 min; linear increase to 95% B over 1 min; constant 95% B for 9 min; linear decrease to 3% B over 5 min. The total running time, including initial conditioning of the column, was 90 min. The eluted peptides were transferred to the nanoelectrospray source of a quadrupole TOF-MS instrument (Synapt High Definition Mass Spectrometry system, Waters) via a Teflon capillary union and pre-cut PicoTip (Waters). ESI was used with a capillary voltage of 3 kV and sampling cone voltage of 35 V. Low (6 eV) or elevated (step from 15 to 30 eV) collision energy was used to generate either intact peptide precursor ions (low energy) or peptide product ions (elevated energy). The source temperature was 100°C, and the detector was operated in the positive-ion mode. Data were collected in the *m*/*z* = 300–2000 range using an independent reference spray and the NanoLockSpray interference procedure in which Glu-1-fibrinopeptide B (*m*/*z* = 785.8426) was infused via the NanoLockSpray ion source and sampled every 10 s for external mass calibration. Data were collected using MassLynx software (v4.1, Waters). ProteinLynx Global Server Browser (v2.3, Waters) was used to identify the protein based on its peptide mass fingerprints.

### Measurement of zinc concentration

Each sample was added to an acid-washed test tube containing nitric acid (0.3 mL) and H_2_O_2_ (0.1 mL) and digested at 130°C for 2 d in an aluminum dry bath block. The evaporated samples were dissolved in deionized distilled water, and zinc concentrations were measured using ICP-MS (ICPMS-2030, Shimadzu, Japan). A ZnSO_4_ solution was used as a concentration standard.

### FT-ICR-MALDI-TOF/MS

Recombinant human Zn_7_GIF/MT-3 was incubated with 0.1 N HCl at 37°C for 30 min and then exchanged with 20 mM Tris-HCl (pH 7.5) buffer at 37°C for 36 h to prepare oxidized apo-GIF/MT-3. Low-molecular-weight molecules were then removed using a 3 kDa Amicon Ultra centrifugal filter unit. Zn_7_GIF/MT-3 or apo-GIF/MT-3 (0.5 µL) in 20 mM Tris-HCl (pH 7.5) were mixed with a solution of α-cyano-4-hydroxycinnamic acid matrix (0.5 µL, 60% v/v acetonitrile, 0.2% v/v trifluoroacetic acid) and then dispensed into 384-well plates. The crystals obtained on the plate were analyzed using FT-ICR-MS (7T solariX, Bruker Daltonics, Billerca, MA, USA) equipped with a MALDI source operating in positive ion mode. Analytical conditions were as follows: *m*/*z* range, 1000–10,000; number of scans averaged, 3; accumulation time, 2.00 s; polarity, positive.

### Raman Spectroscopy

Recombinant Zn_7_GIF/MT-3 was incubated with 0.1 N HCl for 30 min and then replaced with 20 mM Tris-HCl (pH 7.5) buffer for 36 h at 37°C (apo-GIF/MT-3) or HPE-IAM (5 mM) in 20 mM Tris-HCl (pH 7.5) for 36 h at 60°C. Then, low-molecular-weight molecules were removed using a 3 kDa Amicon Ultra centrifugal filter unit. The resulting protein was concentrated to ≈4 mg/mL using a 5 kDa centrifugal concentrator (Vivaspin; Sartorius, Gottingen, Germany). For drop-deposition Raman spectroscopy, 0.5 μL of each protein sample was first dried onto a hydrophobic quartz coverslip for up to 15 min under vacuum. Spectra were then collected from the “coffee ring” of each drop, where proteins were found in the absence of bulk salt, using a Raman microscope system with a charge-coupled-device detector (InVia, Renishaw, New Mills, UK). Each sample was excited using a 785 nm diode laser focused through a Leica 50× (0.75 numerical aperture) short-working-distance air objective, with ≈100 mW power incident on each sample. The laser was focused onto the sample using an on-screen camera. WiRE software (v4.1, Renishaw) was used for spectral acquisition, data collection, and cosmic ray removal. The Raman system was calibrated against the 520 cm^−1^ reference peak of silicon prior to each experiment. All spectra were processed using the IrootLab plugin (0.15.07.09-v) in MATLAB (The MathWorks, Inc., MA, United States). The background was carefully subtracted from the spectra using blank quartz spectra, then the background-corrected spectra were smoothed using a wavelet denoising function. Fluorescence was removed by fitting and subtracting a fifth-order polynomial, and the ends of each spectrum were anchored to the axis using a rubber-band-like function before intensity normalization.

### Detection of protein-bound sulfane sulfur

LC-ESI-MS/MS analysis with HPE-IAM was used to determine the levels of protein-bound sulfane sulfur. Some reagents such as H_2_O_2_, SNAP, TCEP, and KCN were filtered through a 3 kDa Amicon Ultra centrifugal filter unit prior to use. A high-molecular-weight cytosolic fraction from mouse brain or purified MT protein solution was incubated with HPE-IAM under the appropriate conditions to yield bis-S-HPE-AM adducts. Incubation with HPE-IAM at temperatures >60°C was not suitable because it yielded false-positives for sulfane sulfur in protein-free negative controls. The resulting solutions were filtered through a 3 kDa Amicon Ultra centrifugal filter unit to obtain low-molecular-weight fractions containing bis-S-HPE-AM adducts. HPE-AM adducts were diluted four-fold with 0.1% v/v formic acid containing known amounts of isotope-labeled internal standard (bis-S^34^-HPE-AM^6^) and the sulfane sulfur concentrations were determined using LC-ESI-MS/MS. A triple quadrupole mass spectrometer (EVOQ Qube, Bruker) coupled to an ultra-high-performance LC system (Advance, Bruker) was used to perform LC-ESI-MS/MS. Sulfane sulfur-derived bis-S-HPE-AM was separated using a YMC-Triart C_18_ column (50 mm × 2.0 mm i.d.) at 40°C. Mobile phases A (0.1% v/v formic acid) and B (0.1% v/v formic acid in methanol) were linearly mixed at a flow rate of 0.2 mL/min using a gradient system as follows: 3% B for 3 min; linear increase to 95% B over 12 min; constant 95% B for 1 min; linear decrease to 3% B. MS spectra were obtained using a heated ESI source with the following settings: spray voltage, 4000 V; cone temperature, 350°C; heated probe temperature, 250°C; cone gas flow, 25 psi; probe gas flow, 50 psi; nebulizer gas flow, 50 psi.

### Measurement of free SH/SSH content

Ellman’s reagent (DTNB) was used to estimate the concentration of sulfhydryl groups in GIF/MT-3 by comparison with a standard curve of the sulfhydryl-containing compound GSH. Briefly, after removal of low-molecular-weight molecules, GIF/MT-3 protein (1 µM) was incubated with DTNB (500 µM) in 100 mM Tris-HCl (pH 8.0) and 1 mM EDTA at room temperature for 5 min, and the absorbance at 412 nm was measured. A putative reaction scheme for DTNB with RSSH is shown in Fig S8.

### NADPH consumption

To study the kinetics of reduction of human apo-GIF/MT-3 by Trx, 200 µL reaction mixtures containing 100 mM KPi (pH 7.5), 100 µM NADPH, 1 µg human Trx (0.5 µM), 0.7 µg rat TrxB (50 nM), and the concentrations of oxidized apo-GIF/MT-3 indicated in Fig. 7A were used. To study the reduction of human apo-MT-3 or human insulin by Trx, 200 µL reaction mixtures containing 100 mM KPi (pH 7.5), 100 µM NADPH, 1 µg human Trx (0.5 µM), 0.7 µg rat TrxB (50 nM), and the concentrations of oxidized apo-GIF/MT-3 or human insulin indicated in Fig. 7B were used. To compare the reduction of human apo-GIF/MT-3 or Zn_7_GIF/MT-3 by Trx, TRP14, or TRP32, 200 µL reaction mixtures containing 100 mM KPi (pH 7.5), 100 µM NADPH, 6 µg of human Trx, TRP14, or TRP32, 1 µg of rat TrxB, and 6 µg of oxidized apo-GIF/MT-3 or Zn_7_GIF/MT-3 were used (Fig. 7C). Reactions were performed at room temperature and NADPH oxidation was monitored by measuring the absorbance of NADP^+^ at 340 nm. To restore sulfane sulfur in apo-GIF/MT-3 after incubation with the Trx/TR system (Fig. 7D), apo-GIF/MT-3 (5 µM) was incubated with Trx (6 µM), TrxB (72 nM), and NADPH (100 µM) at 25°C for 30 min in 100 mM KPi (pH 7.5) and then with 5 mM HPE-IAM. Sulfane sulfur content in apo-GIF/MT-3 was determined using LC-ESI-MS/MS after 3 kDa filtration, with the peak intensity obtained without apo-GIF/MT-3 being subtracted from that obtained with the complete mixture.

### Homology modeling of MT isoforms

Homology modeling of human GIF/MT-3 was performed using MOE software (2018.01; Chemical Computing Group ULC, 1010 Sherbrooke St. West, Suite #910, Montreal, QC, Canada, H3A 2R7, 2018). To construct a model of GIF/MT-3 (National Center for Biotechnology Information reference sequence: NP_005945.1), the crystal structure of rat MT-2 (PDB code: 4MT2) and the NMR structure of the α domain of human GIF/MT-3 (PDB code: 2F5H) were used as templates. Following the alignment of the primary structures (Fig. S3A), the sequence similarities of the templates were 67.7% and 100% compared with the β domain and α domain of GIF/MT-3, respectively. The metals in the templates were changed as desired or deleted to model apo-GIF/MT-3. To construct the GIF/MT-3 model, 100 independent models of the target protein were built using the segment-matching procedure in MOE. Refinement of the model with the lowest generalized Born/volume integral (GVBI) was achieved by energy minimization of outlier residues in Ramachandran plots generated within MOE. The final model of GIF/MT-3 exhibited a 3D structure similar to those of MT2 and the α domain of GIF/MT-3 (Fig. S3B). The same method was used to construct homology models of human MT-1A and MT-2, with rat MT-2 as the template for MT-1, and both rat MT-2 and the α domain of human MT-2 (PDB code: 1MHU) as templates for MT2 (Fig. S4).

### Generation of sulfane sulfur-bound 3D model of MT

All cysteine residues in the homology model of MT were replaced with cysteine persulfide or polysulfides by performing a residue scan using the Protein Design module of MOE, and the resultant changes in metal binding affinities and complex stabilities were evaluated. The orientation of cysteine persulfide was manually modified to increase its interaction with the metals. Supplementary data file containing homology model of GIF/MT-3 with replacement of all cysteine residues by cysteine persulfide in PDB format (*.pdb) was linked to the article (Data S1).

### Protein thermostability and metal-binding affinity scoring

We assessed the influence of sulfane sulfur on protein unfolding free energy using the Protein Design module of MOE, which computed a stability scoring function, ΔΔGs, based on the GBVI and weighted surface area:

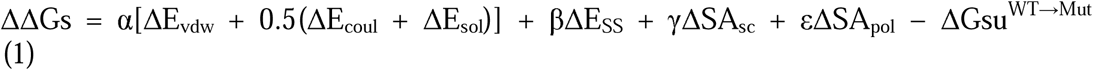

where ΔE_vdw_ is the AMBER van der Waals interaction energy, ΔE_coul_ is the AMBER Coulomb interaction energy, ΔE_sol_ is the change in solvation energy calculated using the GBVI, and E_SS_ is the change in energy due to the presence of a disulfide bond. ΔSAsc and ΔSApol are the changes in the side-chain and polar surface areas, respectively. α is a scaling factor accounting for configurational entropy effects, and ΔGsu^WT→Mut^ is the change in stability of the unfolded states. The affinity score was also calculated using MOE software as the difference between the potential energy values of the protein, free zinc, and metal–protein complex.

### Quantum chemistry calculations

Since the whole structure of the GIF/MT-3 is quite large, we divided GIF/MT-3 into two domains (α-domain and β-domain) in the quantum chemistry calculations in order to assign the observed Raman spectra. We have independently constructed α-domain and β-domain models of apo-GIF/MT-3 with disulfide bonds between neighboring cysteines or tetrasulfide bonds. These models are referred to as apo-GIF/MT-3_S2 and apo-GIF/MT-3_S4, respectively, and are shown in Fig. 1E–H. The initial structures were taken from the results of the homology modeling by MOE (see Fig. 9A). H atoms were placed instead of Zn-S bonds in the models. To consider apo-GIF/MT-3 models, the nearest S atoms are supposed to form disulfide or tetrasulfide bond. The Raman spectra were computed by frequency calculation. All quantum chemical calculations were carried out at B3LYP/6-31G(d) level by GAUSSIAN16 (Revision C.01, Gaussian, Inc., Wallingford CT, 2016). After obtaining the Raman spectra for the α- and β-domains, these spectra are summed to obtain the Raman spectra of the apo-GIF/MT-3 as illustrated in Fig. 1I. The corresponding Zn-binding models were constructed and evaluated the Raman spectra with the same manner.

### Statistical analysis

All reported data represent the mean ± SD of three independent experiments, except for the MOE calculations. The statistical significance of pair-wise differences was assessed using Student’s *t*-test. *p* < 0.05 was considered to indicate a statistically significant difference, and *p* < 0.01 was considered highly significant.

## Supporting information

Response to reviewers

## Acknowledgments

We thank Prof. T. Sawa (Kumamoto University, Japan) for kind donation of NAC derivatives and Prof. E. S. J. Arnér (Karolinska Institutet, Sweden) for kind donation of TRP14 and TRP32.

## Author contributions

Y.Shinkai, Y.D., and Y.K. designed the experiments. Y.Shinkai and Y.D. performed the experiments and analyzed the results, except for the Raman analysis. G.D. and S.M. performed the Raman analysis. T.M. and Y. Shigeta performed the computational analysis. M.A., T.S., M.N., T.I., and T.A. provided useful information and theoretical support. Y.Shinkai wrote the original draft of the manuscript. Y.Shinkai, S.M., J.F., Y.Shigeta, and Y.K. edited the manuscript. Y.K. supervised the study. All authors read and approved the final manuscript.

## Data and materials availability

All data are available in the main text or the supplementary materials.

## Supplementary Materials

Supplementary material for this article is available.

Fig. S1. Zn-binding GIF/MT-3 models and calculated Raman spectra.

Fig. S2. Determination of sulfane sulfur and zinc content in wild-type and several mutant proteins.

Fig. S3. Homology modeling of GIF/MT-3.

Fig. S4. 3D structural models of human Zn_7_MT-1A and Zn_7_MT-2

Fig. S5. Schematic structures of Zn_7_GIF/MT-3 and Zn_7_S_20_GIF/MT-3.

Fig. S6. Zn_3_Cys_9_ and Zn_4_Cys_11_ clusters containing the polysulfide form of sulfane sulfur.

Fig. S7. Amino acid sequences of human MTs and GIF/MT-3 mutant

Fig. S8. A putative reaction scheme for DTNB with RSSH.

Table. S1. Peak assignments for apo-GIF/MT-3 model structures

Table. S2. Peak assignments for Zn_7_S_20_GIF/MT-3 and Zn_7_GIF/MT-3 model structures

Table. S3. Thermostability score of sulfane sulfur-bound MT isoforms with or without Zn.

Table. S4. Fragment sequences of a mouse brain sulfane sulfur-binding protein, determined using nano-UPLC-MS.

Data. S1. Pdb file of Zn_7_S_20_GIF/MT-3 generated by homology modelling.

## References

1. J. I. Toohey, Sulfur signaling: is the agent sulfide or sulfane? Anal. Biochem. 413, 1–7 (2011).

2. M. Iciek, A. Bilska-Wilkosz, M. Gorny, Sulfane sulfur - new findings on an old topic. Acta Biochim. Pol. 66, 533–544 (2019).

3. Y. Shinkai, Y. Kumagai, Sulfane Sulfur in Toxicology: A Novel Defense System Against Electrophilic Stress. Toxicol. Sci. 170, 3–9 (2019).

4. P. D. Kruithof, S. Lunev, S. P. Aguilar Lozano, F. de Assis Batista, Z. M. Al-Dahmani, J. A. Joles, A. M. Dolga, M. R. Groves, H. van Goor, Unraveling the role of thiosulfate sulfurtransferase in metabolic diseases. Biochim Biophys Acta Mol Basis Dis 1866, 165716 (2020).

5. B. Pedre, T. P. Dick, 3-Mercaptopyruvate sulfurtransferase: an enzyme at the crossroads of sulfane sulfur trafficking. Biol. Chem. 402, 223–237 (2021).

6. T. Ida, T. Sawa, H. Ihara, Y. Tsuchiya, Y. Watanabe, Y. Kumagai, M. Suematsu, H. Motohashi, S. Fujii, T. Matsunaga, M. Yamamoto, K. Ono, N. O. Devarie-Baez, M. Xian, J. M. Fukuto, T. Akaike, Reactive cysteine persulfides and S-polythiolation regulate oxidative stress and redox signaling. Proc. Natl. Acad. Sci. U. S. A. 111, 7606–7611 (2014).

7. J. M. Fukuto, L. J. Ignarro, P. Nagy, D. A. Wink, C. G. Kevil, M. Feelisch, M. M. Cortese-Krott, C. L. Bianco, Y. Kumagai, A. J. Hobbs, J. Lin, T. Ida, T. Akaike, Biological hydropersulfides and related polysulfides - a new concept and perspective in redox biology. FEBS Lett. 592, 2140–2152 (2018).

8. T. Akaike, T. Ida, F. Y. Wei, M. Nishida, Y. Kumagai, M. M. Alam, H. Ihara, T. Sawa, T. Matsunaga, S. Kasamatsu, A. Nishimura, M. Morita, K. Tomizawa, A. Nishimura, S. Watanabe, K. Inaba, H. Shima, N. Tanuma, M. Jung, S. Fujii, Y. Watanabe, M. Ohmuraya, P. Nagy, M. Feelisch, J. M. Fukuto, H. Motohashi, Cysteinyl-tRNA synthetase governs cysteine polysulfidation and mitochondrial bioenergetics. Nat Commun 8, 1177 (2017).

9. Y. Abiko, E. Yoshida, I. Ishii, J. M. Fukuto, T. Akaike, Y. Kumagai, Involvement of reactive persulfides in biological bismethylmercury sulfide formation. Chem. Res. Toxicol. 28, 1301–1306 (2015).

10. N. M. Giles, A. B. Watts, G. I. Giles, F. H. Fry, J. A. Littlechild, C. Jacob, Metal and redox modulation of cysteine protein function. Chem. Biol. 10, 677–693 (2003).

11. M. Margoshes, B. L. Vallee, A cadmium protein from equine kidney cortex. J. Am. Chem. Soc. 79, 4813–4814 (1957).

12. Y. J. Kang, Metallothionein redox cycle and function. Exp Biol Med (Maywood) 231, 1459–1467 (2006).

13. W. Maret, Metallothionein redox biology in the cytoprotective and cytotoxic functions of zinc. Exp. Gerontol. 43, 363–369 (2008).

14. A. Krezel, Q. Hao, W. Maret, The zinc/thiolate redox biochemistry of metallothionein and the control of zinc ion fluctuations in cell signaling. Arch. Biochem. Biophys. 463, 188–200 (2007).

15. M. Capdevila, J. Domenech, A. Pagani, L. Tio, L. Villarreal, S. Atrian, Zn- and Cd-metallothionein recombinant species from the most diverse phyla may contain sulfide (S2-) ligands. Angew. Chem. Int. Ed. Engl. 44, 4618-4622 (2005).

16. L. Tio, L. Villarreal, S. Atrian, M. Capdevila, The Zn- and Cd-clusters of recombinant mammalian MT1 and MT4 metallothionein domains include sulfide ligands. Exp. Biol. Med. (Maywood) 231, 1522–1527 (2006).

17. Y. Uchida, K. Takio, K. Titani, Y. Ihara, M. Tomonaga, The growth inhibitory factor that is deficient in the Alzheimer’s disease brain is a 68 amino acid metallothionein-like protein. Neuron 7, 337–347 (1991).

18. A. Torreggiani, A. Tinti, Raman spectroscopy a promising technique for investigations of metallothioneins. Metallomics 2, 246–260 (2010).

19. G. Devitt, K. Howard, A. Mudher, S. Mahajan, Raman Spectroscopy: An Emerging Tool in Neurodegenerative Disease Research and Diagnosis. ACS Chem. Neurosci. 9, 404–420 (2018).

20. R. Steudel, T. Chivers, The role of polysulfide dianions and radical anions in the chemical, physical and biological sciences, including sulfur-based batteries. Chem. Soc. Rev. 48, 3279–3319 (2019).

21. A. W. Schwab, W. K. Rohwedder, J. S. Ard, Mass, nuclear magnetic resonance, and Raman spectroscopy of octadecyl sulfide, disulfide, and tetrasulfide. Phosphorus and Sulfur and the Related Elements 7, 315–319 (1979).

22. T. Chivers, C. Lau, Raman spectroscopic identification of the S4N- and S3- ions in blue solutions of sulfur in liquid ammonia. Inorg. Chem. 21, 453–455 (1982).

23. G. W. Irvine, N. Korkola, M. J. Stillman, Isolated domains of recombinant human apo-metallothionein 1A are folded at neutral pH: a denaturant and heat-induced unfolding study using ESI-MS. Biosci. Rep. 38, (2018).

24. D. A. Capdevila, B. J. C. Walsh, Y. Zhang, C. Dietrich, G. Gonzalez-Gutierrez, D. P. Giedroc, Structural basis for persulfide-sensing specificity in a transcriptional regulator. Nat. Chem. Biol. 17, 65–70 (2021).

25. G. L. Ellman, Tissue sulfhydryl groups. Arch. Biochem. Biophys. 82, 70–77 (1959).

26. W. Maret, The redox biology of redox-inert zinc ions. Free Radic. Biol. Med. 134, 311–326 (2019).

27. W. Maret, B. L. Vallee, Thiolate ligands in metallothionein confer redox activity on zinc clusters. Proc. Natl. Acad. Sci. U. S. A. 95, 3478–3482 (1998).

28. K. Ono, T. Akaike, T. Sawa, Y. Kumagai, D. A. Wink, D. J. Tantillo, A. J. Hobbs, P. Nagy, M. Xian, J. Lin, J. M. Fukuto, Redox chemistry and chemical biology of H2S, hydropersulfides, and derived species: implications of their possible biological activity and utility. Free Radic. Biol. Med. 77, 82–94 (2014).

29. J. Lu, A. Holmgren, The thioredoxin antioxidant system. Free Radic. Biol. Med. 66, 75–87 (2014).

30. A. Holmgren, Reduction of disulfides by thioredoxin. Exceptional reactivity of insulin and suggested functions of thioredoxin in mechanism of hormone action. J. Biol. Chem. 254, 9113–9119 (1979).

31. M. Vasak, Advances in metallothionein structure and functions. J. Trace Elem. Med. Biol. 19, 13–17 (2005).

32. A. Ziller, L. Fraissinet-Tachet, Metallothionein diversity and distribution in the tree of life: a multifunctional protein. Metallomics 10, 1549–1559 (2018).

33. E. Cuevasanta, M. Lange, J. Bonanata, E. L. Coitino, G. Ferrer-Sueta, M. R. Filipovic, B. Alvarez, Reaction of Hydrogen Sulfide with Disulfide and Sulfenic Acid to Form the Strongly Nucleophilic Persulfide. J. Biol. Chem. 290, 26866–26880 (2015).

34. H. A. Hamid, A. Tanaka, T. Ida, A. Nishimura, T. Matsunaga, S. Fujii, M. Morita, T. Sawa, J. M. Fukuto, P. Nagy, R. Tsutsumi, H. Motohashi, H. Ihara, T. Akaike, Polysulfide stabilization by tyrosine and hydroxyphenyl-containing derivatives that is important for a reactive sulfur metabolomics analysis. Redox Biol 21, 101096 (2019).

35. T. Sawa, T. Takata, T. Matsunaga, H. Ihara, H. Motohashi, T. Akaike, Chemical Biology of Reactive Sulfur Species: Hydrolysis-Driven Equilibrium of Polysulfides as a Determinant of Physiological Functions. Antioxid Redox Signal, (2021).

36. M. R. Filipovic, J. Zivanovic, B. Alvarez, R. Banerjee, Chemical Biology of H2S Signaling through Persulfidation. Chem. Rev. 118, 1253–1337 (2018).

37. N. Krishnan, C. Fu, D. J. Pappin, N. K. Tonks, H2S-Induced sulfhydration of the phosphatase PTP1B and its role in the endoplasmic reticulum stress response. Sci Signal 4, ra86 (2011).

38. E. Doka, I. Pader, A. Biro, K. Johansson, Q. Cheng, K. Ballago, J. R. Prigge, D. Pastor-Flores, T. P. Dick, E. E. Schmidt, E. S. Arner, P. Nagy, A novel persulfide detection method reveals protein persulfide- and polysulfide-reducing functions of thioredoxin and glutathione systems. Sci Adv 2, e1500968 (2016).

39. R. Wedmann, C. Onderka, S. Wei, I. A. Szijarto, J. L. Miljkovic, A. Mitrovic, M. Lange, S. Savitsky, P. K. Yadav, R. Torregrossa, E. G. Harrer, T. Harrer, I. Ishii, M. Gollasch, M. E. Wood, E. Galardon, M. Xian, M. Whiteman, R. Banerjee, M. R. Filipovic, Improved tag-switch method reveals that thioredoxin acts as depersulfidase and controls the intracellular levels of protein persulfidation. Chem. Sci. 7, 3414–3426 (2016).

40. Y. Yang, W. Maret, B. L. Vallee, Differential fluorescence labeling of cysteinyl clusters uncovers high tissue levels of thionein. Proc. Natl. Acad. Sci. U. S. A. 98, 5556–5559 (2001).

41. D. Sagher, D. Brunell, J. F. Hejtmancik, M. Kantorow, N. Brot, H. Weissbach, Thionein can serve as a reducing agent for the methionine sulfoxide reductases. Proc. Natl. Acad. Sci. U. S. A. 103, 8656–8661 (2006).

42. M. Vasak, G. Meloni, Mammalian Metallothionein-3: New Functional and Structural Insights. Int. J. Mol. Sci. 18, (2017).

43. Y. Uchida, Molecular mechanisms of regeneration in Alzheimer’s disease brain. Geriatr Gerontol Int 10 **Suppl 1**, S158–168 (2010).

44. J. Y. Koh, S. J. Lee, Metallothionein-3 as a multifunctional player in the control of cellular processes and diseases. Mol. Brain 13, 116 (2020).

45. J. Jia, X. Zeng, G. Xu, Z. Wang, The Potential Roles of Redox Enzymes in Alzheimer’s Disease: Focus on Thioredoxin. ASN Neuro 13, 1759091421994351 (2021).

46. S. Akterin, R. F. Cowburn, A. Miranda-Vizuete, A. Jimenez, N. Bogdanovic, B. Winblad, A. Cedazo-Minguez, Involvement of glutaredoxin-1 and thioredoxin-1 in beta-amyloid toxicity and Alzheimer’s disease. Cell Death Differ. 13, 1454–1465 (2006).

47. C. Andreini, L. Banci, I. Bertini, A. Rosato, Counting the zinc-proteins encoded in the human genome. J. Proteome Res. 5, 196–201 (2006).

48. M. Lange, K. Ok, G. D. Shimberg, B. Bursac, L. Marko, I. Ivanovic-Burmazovic, S. L. J. Michel, M. R. Filipovic, Direct Zinc Finger Protein Persulfidation by H2 S Is Facilitated by Zn(2). Angew. Chem. Int. Ed. Engl. 58, 7997-8001 (2019).

49. K. Zhao, S. Li, L. Wu, C. Lai, G. Yang, Hydrogen sulfide represses androgen receptor transactivation by targeting at the second zinc finger module. J. Biol. Chem. 289, 20824–20835 (2014).

50. A. Dey, S. Prabhudesai, Y. Zhang, G. Rao, K. Thirugnanam, M. N. Hossen, S. K. D. Dwivedi, R. Ramchandran, P. Mukherjee, R. Bhattacharya, Cystathione beta-synthase regulates HIF-1alpha stability through persulfidation of PHD2. Sci Adv 6, (2020).

51. M. Vasak, Criteria of purity for metallothioneins. Methods Enzymol. 205, 44–47 (1991).

