## Supplementary material for "Growth inhibitory factor/metallothionein-3 is a sulfane sulfur-binding protein": Response to reviewers

Reviewer #2 (Public Review):

Comments on revisions:

Although the authors have revealed the sulfane sulfur content in native MT-3, my question, namely, whether canonical MT-1 and MT-2 contained sulfane sulfur after the induction has been left.

The authors argue that the biological significance of sulfane sulfur in MTs lies in its ability to contribute to metal binding affinity, provide a sensing mechanism against oxidative stress, and aid in the regulation of the protein. Due to their biological roles, induced MT-1 and MT-2 could contain sulfane sulfur in their molecules. Thus, I expect the authors to evaluate or explain the sulfane sulfur content in induced MT-1 and MT-2.

**Our response: Thank you for your valuable comments. In this study, we were not able to examine the role of sulfane sulfur in the induced forms of MT-1 and MT-2. However, this topic is undoubtedly important and intriguing; therefore, we will continue to explore it in future studies.**

Reviewer #3 (Public Review):

Comments on revisions:

The revised manuscript is only slightly changed from the original, with the inclusion of a supplementary figure (Fig. S2) and minor changes in the text. The authors did not choose to carry out the quantitative Zn binding experiment (which I really wanted to see), but given the complexities of the experiment, I'll let it go.

Fig. 9: the authors imply in the mechanistic "redox-switch" figure that Trx/TR can not reduce persulfide linkages. A number of groups have shown this to be the case. I recommend modifying the figure legend or text to make this clear to the reader.

**Our response:** **Thank you for your understanding. Regarding the "redox-switch" figure, although some groups have demonstrated the ability of Trx to reduce persulfide moieties, as you pointed out, we have addressed this discrepancy in the Discussion section as follows (lines 357-361): “In contrast, Trx has been proposed to reduce the persulfide moiety of PTP1B (37) and albumin (38, 39). A possible explanation for this discrepancy is that apo-GIF/MT-3-persulfide is rapidly changed into a different conformation that is topologically resistant to Trx reduction. In other words, Trx may** **exhibit substrate specificity.” Additionally, we have inserted the following sentence just before the above discussion to further clarify this point:“This suggests that the persulfide moiety in GIF/MT-3 appears to be relatively stable against Trx reduction.”**
