## Supplemental Materials for "Growth inhibitory factor/metallothionein-3 is a sulfane sulfur-binding protein"

**This PDF file includes:**

Figs. S1 to S8
Tables S1 to S4

**Other Supplementary Materials for this manuscript include the following:**
Data S1

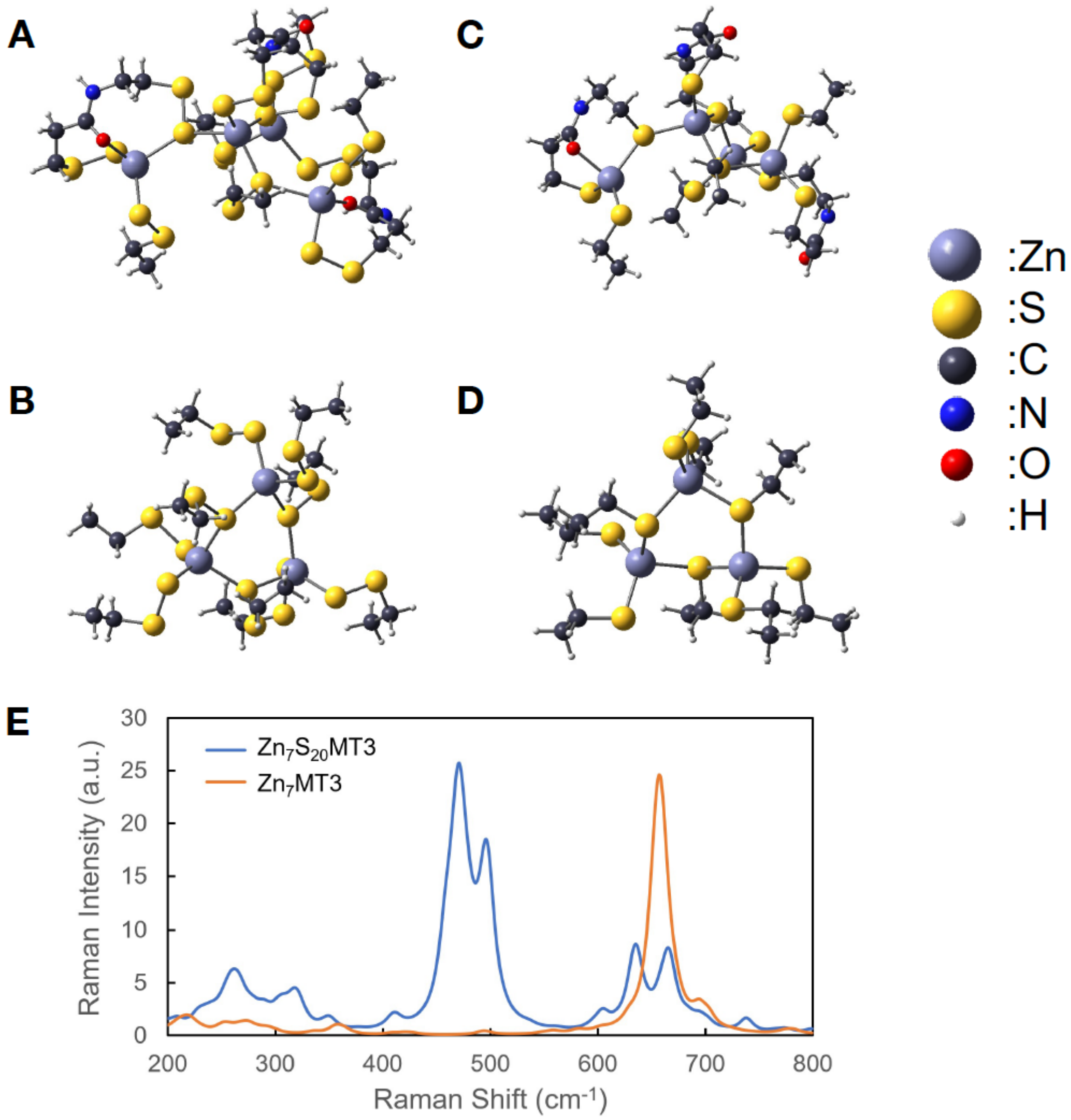


**Fig. S1. Zn-binding GIF/MT3 models and calculated Raman spectra.**
(A) α-domain and (B) β-domain models of $Zn_7S_{20}$GIF/MT3 (assuming all cysteines are persulfides). (C) α-domain and (D) β-domain models of $Zn_7$GIF/MT3 (assuming all cysteines are thiols). (E) Calculated Raman spectra for $Zn_7$GIF/MT3 with or without sulfane sulfurs.

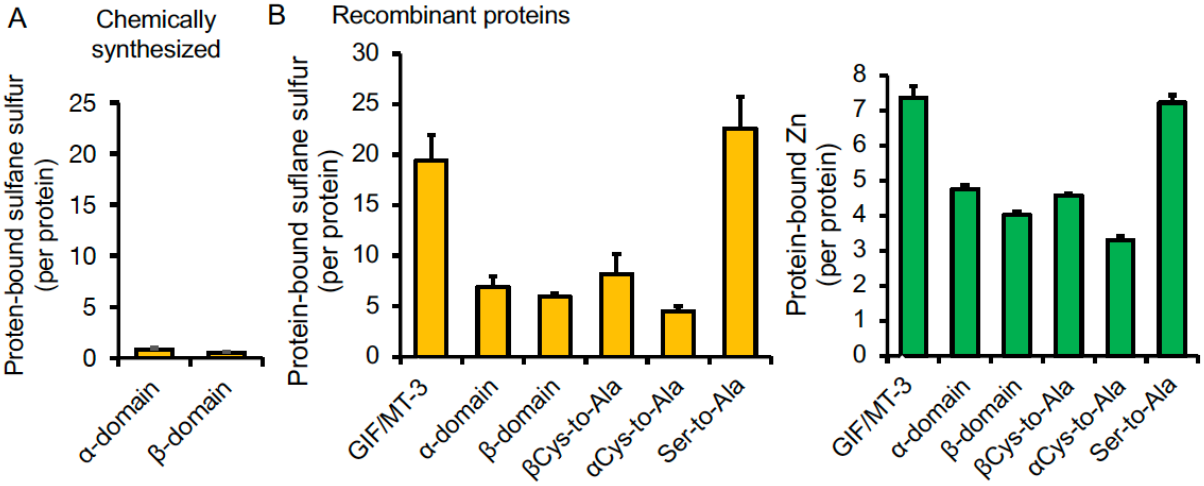


**Fig. S2. Determination of sulfane sulfur and zinc contents of wild-type and several mutant proteins.** (A) Sulfane sulfur detected in chemically synthesized α- and β-domains of GIF/MT-3. (B) Sulfane sulfur (left) and zinc (right) detected in recombinant GIF/MT-3 wild-type and mutant proteins. Sulfane sulfur was quantified using LC–MS after incubation with HPE-IAM (5 mM) at 60°C for 36 h. GIF/MT3-bound zinc content was determined using ICP-MS. The amino acid sequences of GIF/MT-3 mutant proteins are shown in Fig. S7.

**A**

Sequences of MT3 and templates

```
MT3     .MDPETCPCPSGGSCTCADSCKCEGCKCTSCKKSCCSCCPAECEKCAKDCVCKGGEAAEAEAEKCSCCQ
4MT2    XMDP.NCSCATDGSCSCAGSCKCKQCKCTSCKKSCCSCCPVGCAKCSQGCICK.....EA.SDKCSCCA
2F5H    ................................KSCCSCCPAECEKCAKDCVCKGGEAAEAEAEKCSCCQ
```

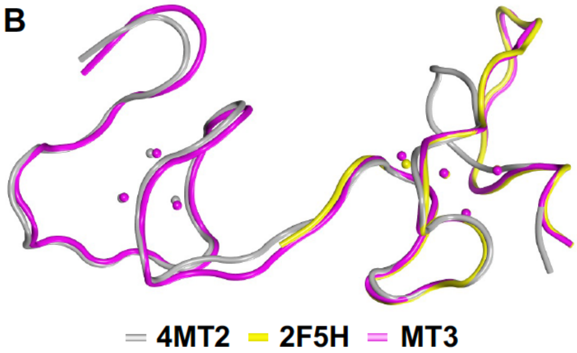


**Fig. S3. Homology modeling of GIF/MT3.**
(A) Sequences of GIF/MT3 and templates. (B) 3D structure of GIF/MT3 and the template structures with PDB codes 4MT2 and 2F5H.

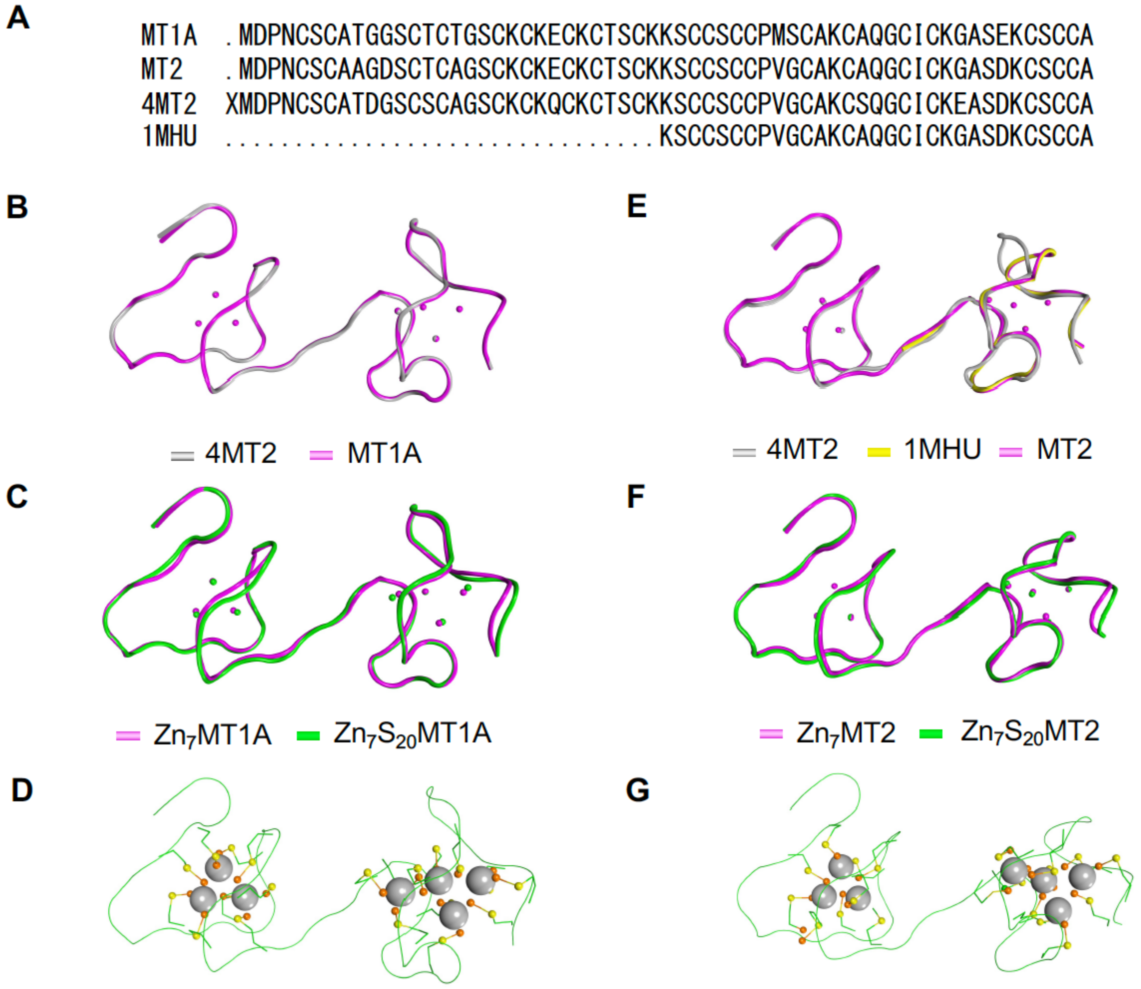


**Fig. S4. 3D structural models of human $Zn_7$MT1A and $Zn_7$MT2.**
(A) Sequences of MT1A and MT2 and the PDB structures with accession numbers 4MT2 and 1MHU. (B) Homology model of $Zn_7$MT1A based on template PDB 4MT2. (C, D) Model 3D structures of $Zn_7S_{20}$MT1A. (E) Homology model of $Zn_7$MT2 based on templates 4MT2 and 1MHU. (F, G) Model 3D structures of $Zn_7S_{20}$MT2.

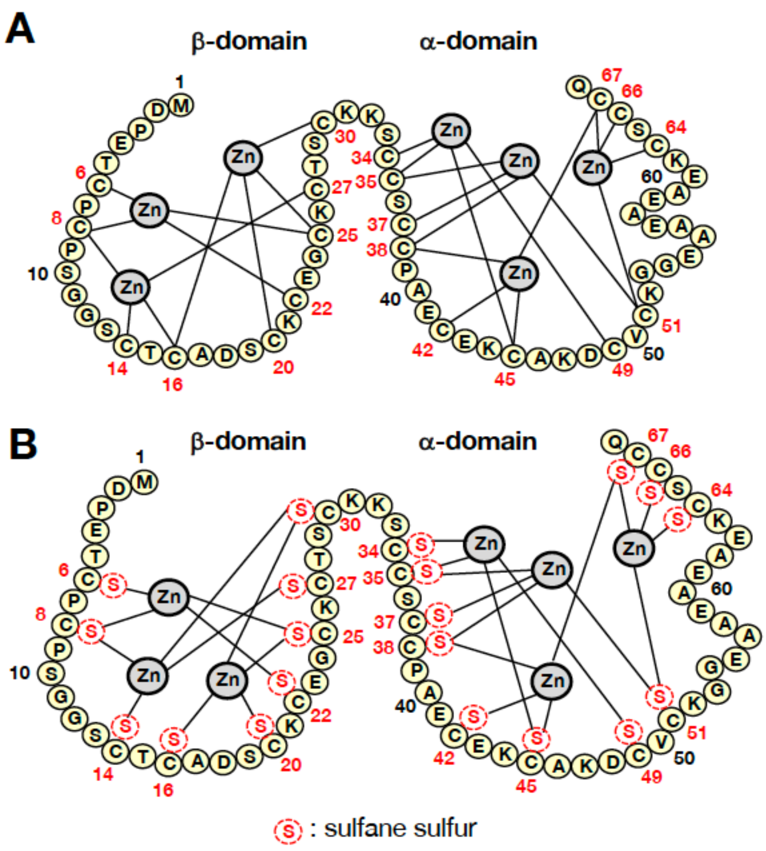


**Fig. S5. Schematic structures of (A) $Zn_7$GIF/MT3 and (B) $Zn_7S_{20}$GIF/MT3.**

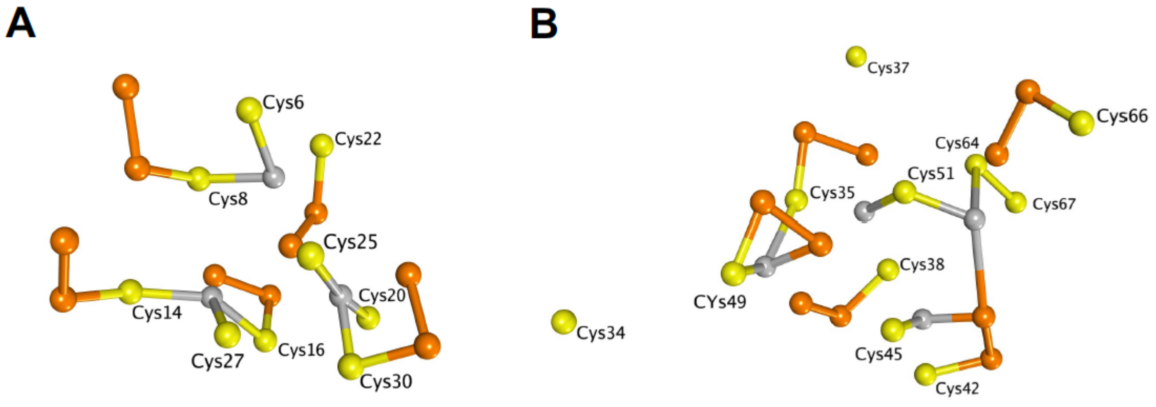


**Fig. S6. $Zn_3Cys_9$ and $Zn_4Cys_{11}$ clusters containing the polysulfide form of sulfane sulfur.** (A) $Zn_3Cys_9$ cluster with 10 sulfane sulfurs (5 RSSSH), taken from the homology model of MT3. (B) $Zn_4Cys_{11}$ cluster with 10 sulfane sulfurs (5 RSSSH), taken from PDB structure 2F5H. Structures were rendered for display using Molecular Operating Environment software. Yellow, orange, and gray spheres indicate cysteine residues, sulfane sulfur atoms, and zinc ions, respectively.

### Human MTs amino acid sequences

| | |
|---|---|
| **MT-1A** | MDPNCSCATG GSCTCTGSCK CKECKCTSCK KSCCSCCPMS CAKCAQGCIC KGASEKCSCC A |
| **MT-2A** | MDPNCSCAAG DSCTCAGSCK CKECKCTSCK KSCCSCCPVG CAKCAQGCIC KGASDKCSCC A |
| **MT-3** | MDPETCPCPS GGSCTCADSC KCEGCKCTSC KKSCCSCCPA ECEKCAKDCV CKGGEAAEAE AEKCSCCQ |

### Recombinant human MTs and their mutant sequences

| | |
|---|---|
| **MT-1A** | GSMDPNCSCATG GSCTCTGSCK CKECKCTSCK KSCCSCCPMS CAKCAQGCIC KGASEKCSCC A |
| **MT-2A** | GSMDPNCSCAAG DSCTCAGSCK CKECKCTSCK KSCCSCCPVG CAKCAQGCIC KGASDKCSCC A |
| **MT-3 wildtype** | GSMDPETCPCPS GGSCTCADSC KCEGCKCTSC KKSCCSCCPA ECEKCAKDCV CKGGEAAEAE AEKCSCCQ |
| **MT-3 Cys-to-Ala** | GSMDPETAPAPS GGSATAADSA KAEGAKATSA KKSAASAAPA EAEKAAKDAV AKGGEAAEAE AEKASAAQ |
| **MT-3 α-domain** | GSCCSCCPA ECEKCAKDCV CKGGEAAEAE AEKCSCCQ |
| **MT-3 β-domain** | GSMDPETCPCPS GGSCTCADSC KCEGCKCTSC KK |
| **MT-3 βCys-to-Ala** | GSMDPETAPAPS GGSATAADSA KAEGAKATSA KKSCCSCCPA ECEKCAKDCV CKGGEAAEAE AEKCSCCQ |
| **MT-3 αCys-to-Ala** | GSMDPETCPCPS GGSCTCADSC KCEGCKCTSC KKSAASAAPA EAEKAAKDAV AKGGEAAEAE AEKASAAQ |
| **MT-3 Ser-to-Ala** | GSMDPETCPCPA GGACTCADAC KCEGCKCTAC KKACCACCPA ECEKCAKDCV CKGGEAAEAE AEKCACCQ |

**Fig. S7. Amino acid sequences of human MTs and GIF/MT-3 mutants.** Mutated amino acids are indicated in red. Amino acids remaining after removal of the GST-tag by thrombin are indicated in blue.

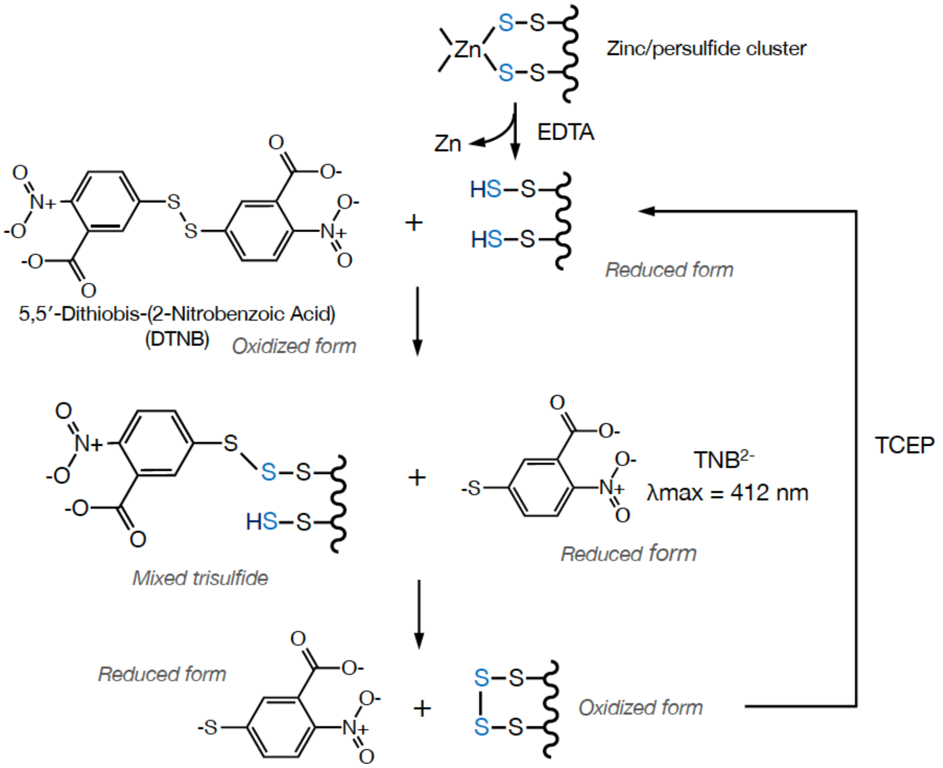


**Fig. S8. A putative reaction scheme for DTNB with RSSH.**

**Table S1.** **Peak assignments for apo-GIF/MT3 model structures.**

| Wavelength | Raman Intensity | Assignment |
|---|---|---|
| | | apo-GIF/MT3_S4 |
| 403.8 | 87.7 | SS-SS (alpha), bent + NH bent |
| 410.6 | 164.0 | C-SS (alpha) bent + NH bent |
| 436.8 | 34.1 | SS-SS(beta), stretch |
| 462.7 | 78.5 | SSS-S (beta), bent |
| 464.2 | 29.7 | SSS-S(beta), stretch |
| 468.6 | 23.9 | SS-SS (alpha), stretch |
| 468.9 | 91.6 | SSS-S (alpha), stretch |
| 480.1 | 18.6 | SSS-S(beta), stretch |
| 482.9 | 45.3 | SS-SS(beta), bent |
| 485.7 | 36.2 | SSS-S (alpha), stretch |
| 489.3 | 12.0 | SSS-S (alpha), stretch |
| 489.6 | 32.3 | S-SH (beta) stretch+ methyl bent |
| 489.6 | 39.2 | S-SH (beta) stretch+ methyl bent |
| 490.8 | 33.2 | S-SH(alpha) stretch + methyl bent |
| 493.3 | 35.4 | S-SH(alpha) stretch + methyl bent |
| 493.6 | 52.5 | S-SH (beta) stretch+ methyl bent |
| 494.2 | 45.7 | S-SH (beta) stretch+ methyl bent |
| 497.8 | 13.9 | SSS-S (alpha), stretch |
| 501.6 | 39.4 | S-SH(alpha) stretch |
| 502.3 | 43.7 | S-SH (beta) stretch |
| 502.3 | 38.1 | S-SH(alpha) stretch |
| 506.1 | 43.6 | S-SH(alpha) stretch |
| 508.9 | 20.9 | SSS-S (alpha), stretch |
| | | apo-GIF/MT3_S2 |
| 391.0 | 12.5 | SS stretch (alpha) + NH stretch |
| 413.8 | 40.6 | SS stretch (beta) |
| 472.3 | 11.0 | NH stretch (alpha) |
| 479.6 | 32.8 | SS stretch (beta) |

**Table S2. Peak assignments for $Zn_7S_{20}$GIF/MT3 and $Zn_7$GIF/MT3 model structures.**

| Wavelength | Raman Intensity | Assignment |
|---|---|---|
| | | $Zn_7S_{20}$GIF/MT3 |
| 469.1 | 37.3 | SS stretch (beta) |
| 469.7 | 56.1 | SS stretch (beta) |
| 471.6 | 41.8 | SS stretch (beta) |
| 472.4 | 57.9 | SS stretch (beta) |
| 473.7 | 71.1 | SS stretch (beta) |
| 478.6 | 69.6 | SS stretch (beta) |
| 493.6 | 30.7 | SS stretch (beta) |
| 494.9 | 117.4 | SS stretch (beta) |
| 498.2 | 137.2 | SS stretch (beta) |
| 408.4 | 20.2 | CH3 torsion, peptide(alpha) |
| 412.5 | 18.4 | CH3 torsion, peptide(alpha) |
| 458.0 | 139.4 | SS stretch (alpha) |
| 462.4 | 39.7 | SS stretch (alpha) |
| 468.0 | 146.7 | SS stretch (alpha) |
| 468.1 | 33.6 | SS stretch (alpha) |
| 471.3 | 79.2 | SS stretch (alpha) |
| 471.7 | 66.4 | SS stretch (alpha) + NH stretch |
| 477.9 | 26.9 | SS stretch (alpha) + NH stretch |
| 486.1 | 34.2 | SS stretch (alpha) + CH3 torsion, peptide |
| 488.3 | 7.2 | CH3 torsion (alpha), peptide |
| 494.9 | 36.7 | SS stretch (alpha) |
| 497.2 | 48.5 | SS stretch (alpha) |
| 497.4 | 34.7 | SS stretch (alpha) |
| 499.1 | 32.8 | NH stretch (alpha) |
| 515.6 | 2.4 | CH3 torsion, peptide(alpha) |
| | | $Zn_7$GIF/MT3 |
| 408.2881 | 4.0012 | CH3 torsion, peptide(alpha) |
| 419.7967 | 4.1649 | CH3 torsion, peptide(alpha) |
| 427.6395 | 3.8385 | CH3 torsion, peptide(alpha) |
| 491.5289 | 0.7488 | CH3 torsion, peptide(alpha) |
| 492.0817 | 3.461 | CH3 torsion, peptide(alpha) |
| 495.6339 | 6.3352 | CH3 torsion, peptide(alpha) |

**Table S3. Thermostability score of sulfane sulfur-bound MT isoforms with or without Zn.** Values were calculated using the Protein Design module in MOE. ΔStability indicates the energy difference between MT with and without sulfane sulfur.

| | Sulfane sulfur | Stability (kcal/mol) | ΔStability (kcal/mol) |
|---|---|---|---|
| $Zn_7$MT1 | | -261 | |
| $Zn_7$MT2 | | -259 | |
| $Zn_7$GIF/MT3 | | -302 | |
| $Zn_7$MT1 | 20 | -368 | -107 |
| $Zn_7$MT2 | 20 | -365 | -106 |
| $Zn_7$GIF/MT3 | 20 | -409 | -107 |
| apo-MT1 | | -190 | |
| apo-MT2 | | -191 | |
| apo-GIF/MT3 | | -221 | |
| apo-MT1 | 20 | -148 | 42 |
| apo-MT2 | 20 | -150 | 41 |
| apo-GIF/MT3 | 20 | -179 | 42 |

**Table S4. Fragment sequences of a mouse brain sulfane sulfur-binding protein, determined using nano-UPLC-MS.**

| Position | Observed MS (Da) | Calculated MS (Da) | Sequence |
|---|---|---|---|
| 2-19 | 2028.75 | 2028.74 | DPETCPCPTGGSCTCSDK + 4 Carbamidomethyl C |
| 33-44 | 1475.48 | 1475.48 | SCCSCCPAGCEK + 5 Carbamidomethyl C |
| 48-63 | 1780.78 | 1780.78 | DCVCKGEEGAKAEAEK + 2 Carbamidomethyl C |

**Data S1. Pdb file of $Zn_7S_{20}$GIF/MT3 generated by homology modelling.**

(separate file)
